# Cross-species insights into placental evolution and diseases at the single-cell resolution

**DOI:** 10.64898/2025.12.26.696571

**Authors:** Guanghui Tan, Ao Zhang, Xuesha Cao, Jingyu Yang, Youjie Cui, Fei Wang, Tao Shi, Hengkuan Li, Haoping Wang, Huiquan Shan, Jilong Ren, Yaqi Zhou, Menghan Wang, Funong Luo, Xi Guo, Wuqiang Huo, Yingran Liu, Zhannur Niyazbekova, Xihong Wang, Zhenyu Xiao, Yi Zheng, Yu Jiang

## Abstract

The placenta is a highly specialized organ in mammals, mediating the exchange of nutrients, gases, and waste between the mother and the fetus while orchestrating intricate immunological interactions to sustain a successful pregnancy. Despite its essential roles, the molecular evolution underlying the diversity of placentas across mammalian species remains largely elusive. Here, we constructed a comprehensive mammalian placental single-cell transcriptomic atlas from approximately 300,000 cells spanning ten species that could well represent the four primary placental types: (discoid, cotyledonary, diffuse, and zonary. Our cross-species analysis highlights trophoblast lineages as key drivers for placental evolution. By reconstructing differentiation trajectories, we elucidate the gene expression dynamics and regulatory networks shaping trophoblast development across diverse placental architectures. Besides, we propose that the association of human trophoblasts with conditions such as pre-eclampsia and miscarriage arises from their unique gene expression profile, which distinguishes them from trophoblasts of other species. The functional experiments further demonstrate that TGIF1 acts as an upstream regulator of key functional genes in extravillous trophoblast cells, modulating their growth, invasion, and migration. Additionally, TGIF1, along with its target genes, such as *ADAM12*, *WNT3A*, and *ZNF831*, is associated with preeclampsia and pregnancy loss. Collectively, these findings provide a high-resolution framework to understand the molecular evolution of the placenta and its role in reproductive success and diseases.

## Introduction

Over a hundred million years of evolution and adaptation, placental mammals have evolved into the vertebrate group with the greatest morphological and genomic diversity^1, 2^. This evolutionary radiation makes the eutherian placenta one of the most morphologically diverse organs in the animal realm^3^. The variation of placentas could typically be ascribed to the interaction between genetically distinct maternal and fetal tissues during placentation^4^. Based on their morphology, eutherian placentas are categorized into four major types based on their morphology: discoid / bidiscoid (e.g., humans, non-human primates, and rodents), zonary (e.g., dogs, cats, and other carnivores), diffuse (e.g., pigs and horses), and multicotyledonary (e.g., cattle and other ruminants) placentas^5^. Among these, the discoid / bidiscoid placenta, also known as the hemochorial placenta, represents the most intimate maternal-fetal interface (interstitial implantation entails the embryo penetrating deep into the uterus and becoming fully engulfed in the endometrial tissue). In this structure, the maternal epithelium and endothelium regress, leaving only the fetal trophoblast and endothelial cells as the barrier between fetal and maternal circulations^6, 7^. Hemochorial placentas can be further classified into three subtypes based on the number of trophoblast layers surrounding the maternal blood: hemomonochorial (primates), hemodichorial (rabbits), and hemotrichorial (rats and mice), with one, two, and three trophoblast layers, respectively. In contrast, the zonary placenta is characterized by degeneration of the maternal uterine epithelium and connective tissues post-implantation, enabling trophoblasts to come into direct contact with the maternal endometrium. The diffuse placenta is characterized by superficial implantation compared to other placental types, with minimal invasion into the uterine lining, and trophoblasts are loosely organized and remain adjacent to the maternal endometrial epithelium without significant tissue destruction or penetration. The final multicotyledonary placenta involves the fusion of specific trophoblasts (binucleate) with a single uterine epithelial cell, leading to trinucleate or even multinucleate structures that include both fetal and maternal components. In general, the diversity in morphology and cell types not only reflects the intricate evolutionary history of placentas but also reveals the distinct evolutionary pathways that different species have followed to meet their specific physiological needs. Indeed, while the morphology of the mammalian placentas has been thoroughly documented in several species^8^, the diversity of their molecular landscape, from the genetic point of view, remains largely enigmatic^9^. In particular, previous studies in some species (e.g., *Homo sapiens*, *Mus musculus*, *Bos taurus*, and *Loxodonta africana*) have revealed significant genetic differences in the placentas, especially during the late stages of pregnancy^9, 10, 11, 12^, that play crucial roles in the crosstalk between placentas and the maternal milieu, but the molecular factors driving the diversification of the placentas during the radiation of mammals remain to be systematically investigated.

The remarkable evolutionary diversity of the mammalian placenta is largely shaped by intraspecific evolutionary conflicts between the mother and the fetus. These conflicts are driven by the dynamic interplay between the maternal-fetal resource allocation and cell-cell interaction. This evolutionary tension not only underpins the extraordinary morphological and functional diversity across species but also contributes to the susceptibility to pregnancy disorders, such as pre-eclampsia (PE), placental accrete spectrum, and recurrent miscarriage, which are disproportionately prevalent in humans^13^. Despite this, the mechanisms underlying placental diversity and the links to many catastrophic complications, especially PE and recurrent pregnancy loss, the incidence of which is high in humans, remain poorly understood. PE, a vascular pregnancy disorder that affects 3-5% of pregnancy^14, 15^, is characterized by insufficient trophoblast invasion and impaired placental perfusion, acting as a catalyst for hypertension and organ damage in the mother, which can, in severe cases, pose a threat to the life of the mother and the fetus^16, 17^. Because of this, PE has garnered mounting attention from both clinicians and the public. Indeed, genetic factors have provided some insights into the etiology of PE^18, 19^, but little is known about the roles of evolutionary adaptation in this reproductive disorder. Recurrent pregnancy loss, defined as two or more consecutive miscarriages with the same partner, occurs in approximately 5% of all pregnancies and represents a significant challenge to successful pregnancy^19, 20^. Despite progress in identification of the contributing factors, such as infections, chromosomal abnormalities, endocrine and metabolic dysfunctions, the antiphospholipid syndrome, as well as uterine anatomical abnormalities, the etiology in approximately half cases remains unknown^21, 22^. This uncertainty hinders treatment and imposes psychological burdens on affected couples.

Cross-species assessment of gene conservation and divergence could offer valuable insights into the mechanisms underlying placental evolution and the associated pregnancy disorders, which is also critical for identification of suitable animal models to investigate human pregnancy-related diseases, thereby advancing translational research^23, 24^. The advancement of high-throughput sequencing techniques such as single-cell RNA-sequencing (scRNA-seq) and single-nucleus RNA-sequencing (snRNA-seq) has greatly facilitated studies aiming to explore the extent to which cell types, the fundamental units of complex tissues, are conserved and divergent across different species^24^, and has also enabled the precise analysis of the heterogeneity and molecular characteristics of different cell types within placentas^25, 26^. For instance, by way of scRNA-seq, the cellular composition of the maternal-fetal interface and the dynamics of trophoblast differentiation in the human placenta^27, 28, 29, 30^, and the conserved features of interspecies placental formation in the placenta of non-human primate cynomolgus macaques (*Macaca fascicularis*)^31^, have been uncovered. The snRNA-seq technology has also been leveraged to unravel the transcriptional profiling of a wide array of cell types and their functions in the mouse and rat placentas^32, 33, 34^. Recently, our spatial transcriptomic analysis of the bovine maternal-fetal interface further advanced the understanding of molecular and cellular interactions in the cotyledonary placentas^35^. Despite all these, current studies remain restricted to individual species, with only a few species covered and focuses on rodents and primates to date, which hinders in-depth understandings of cell biology, evolution, and developmental processes in mammalian placentas. In this sense, it would be beneficial to obtain the cross-species placental molecular data to select appropriate animal models for the enhanced translational relevance, thereby benefiting the clinics in the long run.

In this study, we have for the first time, to our knowledge, constructed a comprehensive single-cell transcriptomic atlas of the mammalian maternal-fetal interface from ten representative species at late pregnancy, enabling unprecedented comparative cross-species analyses of the placental cellular landscape. Our results revealed that the four major cell types (trophoblasts, stromal cells, endothelial cells, and immune cells) showed a gradual increase in the transcriptomic variability among species in relation to evolutionary divergence, with the trophoblast lineage demonstrating the most pronounced variability. We further revealed the transcription factor (TF) regulatory network driving trophoblast evolution. Our findings also elucidated the molecular basis underlying the limited maternal endothelium invasion and the restricted decidualization in the zonary placentas. Finally, we illuminated the evolutionary trajectory of PE, a pregnancy disorder unique to humans, and integrated the genome-wide association study (GWAS) data on pregnancy disorders with single-cell transcriptomic data to identify key cell types, genes, and signaling pathways implicated in miscarriage. Altogether, these findings not only provide novel insights into placental evolution and functions, but also lay the robust foundation for future clinical and translational research that aims to address pregnancy disorders and to improve reproductive health in the human race.

## Results

### Decoding the placental cellular landscape across four placental types in ten mammalian species

To investigate the evolutionary and developmental dynamics of the mammalian placentas at the maternal-fetal interface, we systematically aligned placental samples from ten mammalian species to homologous stages of human gestation, spanning the first, second, and third trimesters based on embryonic developmental timelines and maternal hormonal and physiological milestones (Supplementary Fig.1A and Supplementary Data 1)^36, 37, 38, 39^. To pinpoint the conserved and species-specific features across comparable stages of placental functions, we focused on the late stages of placental development, corresponding to the establishment of definitive placentas, as these represent the period when placental differentiation and functional maturation are largely finalized, while physiological changes associated with the onset of labor have yet occurred (Supplementary Data 2). Specifically, we generated snRNA-seq data from six mammalian species, representing the four major placental types in mammalian species: guinea pigs (*Cavia porcellus*) on embryonic day (E)45.5 of 68, rabbits (*Oryctolagus cuniculus*) on E25.5 of 31-33, dogs (*Canis lupus familiaris*) on E50.5 of 63, cows (*Bos taurus*) on E240 of 284, goats (*Capra hircus*) on E120-145 of 150, and pigs (*Sus scrofa*) on E100 of 114 (Fig.1A-B). To maximize the cross-species comparability while capturing the diversity of placental morphology and functions, we further reanalyzed publicly available scRNA-seq or snRNA-seq data from placentas of four species at the third trimester of pregnancy (Supplementary Fig.1B-E), and the exact developmental timepoints for placental samples from each species were as follows: E266-280 of 280 for humans (*Homo sapiens*)^40^, E140 of 165 for cynomolgus macaques (*Macaca fascicularis*, hereafter referred to as ‘macacas’)^31^, E14.5 of 19-21 for mice (*Mus musculus*)^34^, and E19.5 of 21-23 for rats (*Rattus norvegicus*)^32^. Overall, we compiled a comprehensive dataset of single-cell / -nucleus transcriptomes from 33 libraries, including 19 newly generated datasets from this study, with 1-5 biological replicates per species (Supplementary Fig.1F-K). By using a standardized computational pipeline, data were processed to remove doublets, normalized, corrected for batch effects, reduced in dimension, and clustered separately for each species. Overall, approximately 300,000 high-quality single-cell / -nucleus transcriptomes were obtained for ten species. Cells were annotated into 89 transcriptionally distinct clusters across these ten species, with each characterized by cluster-specific markers (Fig.1C-E).

**Fig.1:**
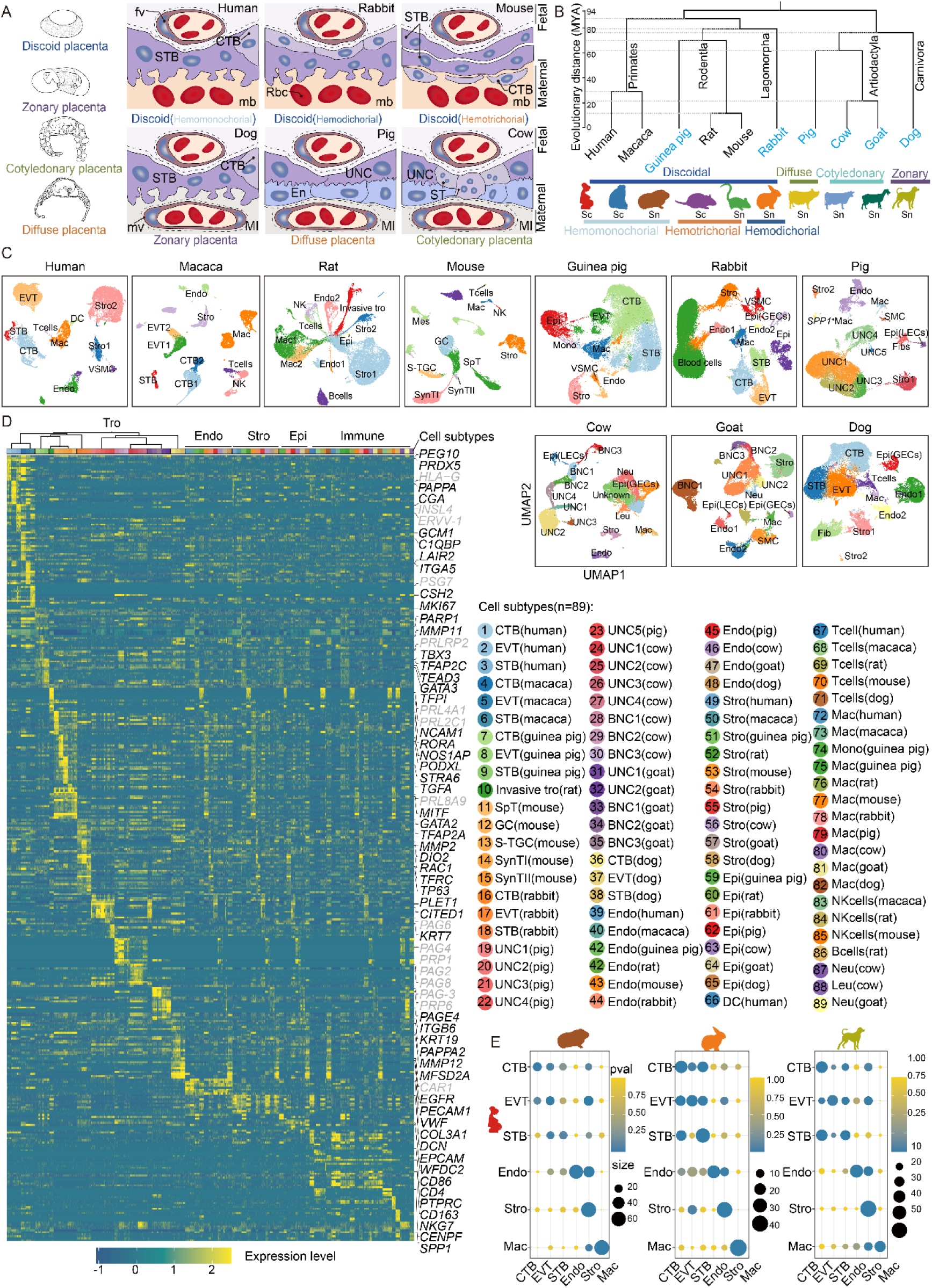
Single-cell / -nucleus transcriptomic mapping and cell type analysis of the mammalian placentas. (A and B) Left: Schematic representation of different types of placentas. The structures and maternal-fetal interfaces of discoid, zonary, cotyledonary, and diffuse placentas are shown. FB, fetal blood; fv, fetal vessel; Ml. maternal interstitium: mv, maternal vessel; ST, specific trophoblast; MB, maternal blood; En, endometrium; Rbc, red blood cell. Right: Phylogeny of the ten mammalian species was analyzed in this work. The scale bar indicates the estimated divergence time (generated via https://phylot.biobyte.de/). MYA, million years ago. Species names in blue indicate data generated in this study, while species names in black represent those reported by previous studies. “Sc” indicates scRNA-seq, and “Sn” represents snRNA-seq. (C) UMAP plots showing clustering results of single-cell or single-nucleus transcriptomes from the placenta or maternal-fetal interface of ten mammalian species: humans^40^, macacas^31^, rats^32^, mice^34^, guinea pigs, rabbits, pigs, cows, goats, and dogs. Each color represents a distinct cell population, including cytotrophoblasts (CTBs), extravillous trophoblasts (EVTs), sncytiotrophoblasts (STB), uninucleate cells (UNCs), binucleate cells (BNCs), stromal cells (Stro, including stromal, fibroblast, and smooth muscle cell subtypes), endothelial cells (Endo), macrophages (Mac), epithelial cells (e.g., Epi, Epi / LEC / GECs comprising luminal and glandular epithelial cells), dendritic (DC), and invasive trophoblasts (Invasive tro). The UMAP plots for biological replicates of each species are shown in Supplementary Fig.1B-K. (D) A heatmap showing the scaled expression levels of cell type-specific markers (left), with each cell type downsampled to 200 cells / nucleus. The columns are color-coded to represent different cell subtypes. Blood cell types were excluded from the analysis, and glandular epithelial cells (GECs) and luminal epithelial cells (LECs) were merged into a single category labeled “Epi”. In addition, certain subclusters were combined into broader categories for clarity, such as Endo1 and Endo2 merged into “Endo”, and Stro1 and Stro2 merged into “Stro”. Expression patterns of 89 representative markers are shown on the right, with more detailed information on marker expression for each cell type provided in Supplementary Fig.1F-K and Supplementary Data 3. Grey genes indicate the missing data owing to the absence of the corresponding orthologue in the species annotation. (E) Dot plots showing the overlap between cell markers for trophoblasts (Tro), stromal cells, as well as endothelial cells reported by previous human scRNA-seq studies and the cluster markers used in the present study. The size of the dots corresponds to the overlap between the cluster gene sets in this study and the gene sets in previous human studies, whereas the color of the dot represents the adjusted *P* value of the one-sided overrepresentation Fisher’s exact test.

We identified a range of trophoblast types, including cytotrophoblasts (CTBs), extravillous trophoblasts (EVTs), and sncytiotrophoblasts (STB) in discoid and zonary placentas, as well as uninucleate cells (UNCs) and binucleate cells (BNCs) in cotyledonary and diffuse placentas (Fig.1C-F and Supplementary Fig.1F-K). Significant overlap of cell types identified in this study with those reported in previous scRNA-seq studies in humans^27, 29, 40^ further validated our annotation (Fig.1F and Supplementary Data 3). The trophoblast subtypes, i.e., CTBs, EVTs, and STB, also exhibited species-specific gene expression patterns. CTBs were annotated by markers such as *ITGB6*, *CDH1*, *MKI67*, *GATA2*, *TFAP2A*, and *PEG10*, consistent with their role as progenitors during placentation^27, 28, 29^. Notably, *KRT7*, previously identified as a pan-trophoblast marker^41^, was extensively expressed across all trophoblast subtypes in species with diffuse and cotyledonary placentas, such as cows, goats, and pigs, corroborating its conserved role in trophoblast identity. EVTs, the invasive trophoblast lineage, exhibited species-specific gene expression, such as *TFPI*, *PRLRP2*, *KRT7*, and *MMP11* in guinea pigs^28, 29, 42, 43^, *MMP2*, *SERPINE1*, and *DIO2* in rabbits^28, 29, 42^, and *PAPPA2*, *SERPINE2*, and *MMP12* in dogs^28, 40^. Notably, the EVTs from guinea pigs specifically expressed the prolactin-related gene (*PRLRP2*), and it was also uniquely and abundantly expressed in the invasive trophoblast lineages of rats and mice^32, 34^, highlighting its conserved role in rodent trophoblast invasion. Rabbit EVTs expressed *NGFR* and *IL6ST*, whilst their canine counterparts expressed *JPT1* and *ANGPT2*. *NOTUM*, a WNT signaling inhibitor, promotes human EVT differentiation by suppressing the canonical WNT signaling^43^, and we observed that *NOTUM* showed specific expression in the EVTs of humans, guinea pigs, and dogs but lower expression in rabbit trophoblasts. *CTNNB1*, which suppresses EVT differentiation upon accumulation^43^, was predominantly expressed in trophoblast stem cells (CTBs) of humans, rabbits, and macacas, while its expression was lower in EVTs. Notably, *CTNNB1* was highly expressed in the EVTs of guinea pigs and dogs, highlighting the evolutionary adaptation of WNT signaling in EVT fate determination across species. Similar to that in humans, markers of deep EVT invasion (*SERPINE1* and *SERPINE2*)^28^ were highly expressed in rabbit EVTs that also expressed human invasive markers (*RAC1*, *ITGA1*, *MMP2*, and *DIO2*). In contrast, canine EVTs highly expressed invasive markers (*MMP12*, *PAPPA2*, *MMP2*, and *ITGA1*), as well as the deep EVT invasion marker *SERPINE2*, but expressed *SERPINE1*, another deep invasive marker, at the lower level (Supplementary Fig.1L), highlighting the differential expression of EVT markers across species. STB also exhibited both interspecies conservation and variability in marker expression, such as *TBX3*, *GATA3*, *TEAD3*, and *MFSD2A* in guinea pigs^27, 28, 29, 44^, *TFRC*, *DUSP9*, and *TP63* in rabbits^31, 45^, and *MFSD2A*, *PTHLH*, and *syncytin-CAR1* in dogs (Supplementary Fig.1H and Supplementary Data 3). Interestingly, *TBX3*, a key TF crucial to trophoblast fusion, was consistently expressed across STB in all examined species, underscoring its fundamental role in placental development and highlighting its evolutionary conservation^44^. Species-specific markers for STB were also identified, such as *NEB* in guinea pigs, *GRB10* in rabbits, and *PHLDB2* and *INHBA* in dogs, reflecting adaptation to the diverse reproductive and physiological demands across species. Notably, some genes exhibited the expression overlap across cell types. For instance, *ADM* was specifically expressed in canine EVTs but highly expressed in both human EVTs and STB; *GGH* was specifically expressed in canine STB but also significantly expressed in human CTBs and STB; *EPAS1* showed specific expression in guinea pig EVTs and canine STB but was extensively expressed in human EVTs and STB (Supplementary Fig.1L). These intricate marker expression patterns reveal the specificity and conservation of placental cell types across species, providing critical molecular evidence to understand the adaptive evolution of the maternal-fetal interface.

We found that in ruminants (cows and goats), UNCs expressed pregnancy-associated glycoproteins (PAGs) such as *PAG2* and *PAG8*, alongside prolactin-related proteins (PRPs) including *PRP1* and *PRP2*. BNCs exhibited high expression of *PAG3*, *PAG4*, *PAG16*, and *PAG17* (Supplementary Fig.1I-J). Additionally, we observed the specific expression of *PAG6* in diffuse placental trophoblasts (pigs) (Supplementary Fig.1K), whereas in multicotyledonary placentas (cows and goats), it was predominantly expressed in BNCs. These markers represent a significantly expanded gene family confined to hooved (ungulate) mammals, reflecting their specialized placental physiology and functions^35, 46^. Trophoblasts in diffuse placentas (e.g., pigs) lacked a syncytialized phenotype and were predominantly represented by UNC subtypes that specifically expressed *PLET1*, *KRT7*, *PPARG*, and *CITED1*^41, 43, 47^.

In addition to the trophoblast lineage, other major cell types, including stromal (Stro), endothelial (Endo), macrophage (Mac), and epithelial (Epi) cells were also annotated, based on the expression of well-established cell type-specific markers (Supplementary Fig.1F-K). The markers expressed in macrophages, stromal cells, and endothelial cells from the placentas of guinea pigs, rabbits, and dogs, which were identified using orthologous genes, also showed significant overlap with those identified in humans (Fisher’s exact test for overrepresentation, *P* < 0.01, Fig.1F). Non-trophoblast cell types also exhibited specific marker expression patterns, such as *DCN* and *COL1A1* in stromal cells, *VWF* and *KDR* in endothelial cells, *CD14* and *C1QA* in macrophages, and *EPCAM* and *WFDC2* in epithelial cells. To elucidate the maternal or fetal origin of these cell types, we further analyzed genetic variants within the snRNA-seq reads using Souporcell, and the result is shown in Supplementary Fig.1M.

### Conserved and divergent cell types during placental evolution

To elucidate the evolutionary conservation and divergence of placental cell types across species, we integrated placental transcriptomic data from the ten mammalian species by using Seurat (v4.4.0)^48^. The results showed that cells and nuclei from different mammalian species were well blended, suggesting high conservation of cell types across species (Fig.2A-B). To delve into the evolutionary relationships among cell types, we employed the pairwise unsupervised MetaNeighbor analysis at the pseudo-cell level, which enables to compare conserved versus divergent cell types across species. By way of this, we identified highly conserved evolutionary features in macrophages, endothelial and stromal cells, while trophoblasts exhibited significantly greater divergence in their evolutionary trajectory (Fig.2C). Next, we analyzed the expression of cell type-specific markers previously validated in mice and primates^27, 30, 31, 33, 34^. These markers were found to be highly expressed in the same placental cell type but barely expressed in other placental cell types, and many of these markers also exhibited similar expression patterns in mammalian species other than mice and primates (Fig.2D). With these markers, we assigned cells within each species to four classes (excluding the undersampled epithelial cells, T cells, dendritic cells, natural killer cells, and B cells): trophoblasts, endothelial cells, stromal cells, and macrophages (Fig.2E). Conserved markers for trophoblasts included critical lineage-determining TFs such as *GATA2*, *GATA3*, and *TFAP2A*^49, 50, 51^ (Fig.2F and Supplementary Fig.2A-B), underscoring the evolutionary importance of these regulators in placental development and trophoblast fate specification.

**Fig.2:**
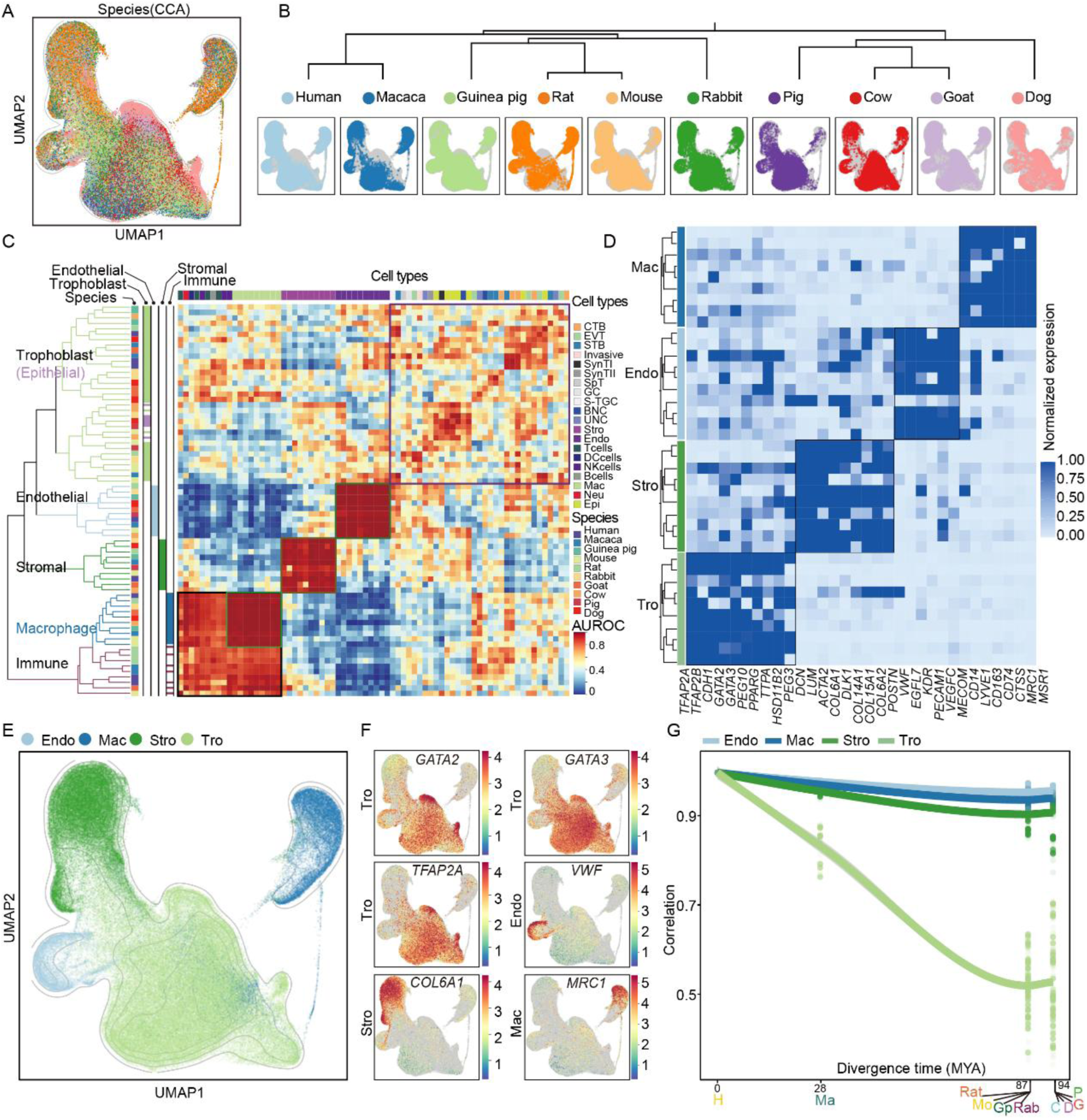
Conserved and divergent cell types during placental evolution. (A) UMAP visualization of integrated transcriptomic data from the placental maternal-fetal interface across multiple species, showing the clustering of different species. Each color represents a species: humans, macacas, guinea pigs, rats, mice, rabbits, pigs, cows, goats, and dogs. (B) A phylogenetic tree depicting the evolutionary relationships among the analyzed species, with corresponding UMAP plots for each species showing the distribution of cell types. The branch length of an evolutionary tree has no meaning. (C) A heatmap illustrating the similarity (AUROC score) among cell types across species. The dendrogram on the left indicates the clustering of cell types based on their AUROC score matrix. (D) A heatmap showing average expression of markers (columns) within each major cell class in ten species (rows). Rows are grouped by cell class (left). Within each class, species are ordered as in Fig.2B. Colors indicating cell classes are uniform in Fig.2C-G. (E) UMAP embedding of integrated cross-species data, with points indicating class identity. (F) Expression levels of subclass-specific markers. *GATA2*, *GATA3*, and *TFAP2A*, markers for trophoblasts, were also expressed in some other cells. *VWF*, a marker for endothelial cells; *COL6A1*, a marker for stromal cells; *MRC1*, a marker for macrophages. Details of gene expression by species are shown in Supplementary Fig.2B. (G) Spearman’s correlations between humans and other species from 100 bootstrap replicates (dots). MYA, million years ago. The evolutionary distances between other species and humans were obtained from Timetree (https://timetree.org).

We further evaluated the interspecies similarity among species classes by analyzing ‘pseudobulk’ transcriptomic profiles on the basis of shared orthologous genes (specified in Methods). Consistent with the results of MetaNeighbor, the cross-correlation analysis of the ten mammalian species revealed that the transcriptomic similarity was primarily driven by cell class identity rather than species identity. For instance, trophoblasts in one species were more closely related to those in other species than they were to other cell classes within the same species. Thus, in the transcriptional profile of placental cells, cell class identity prevailed over species identity (Supplementary Fig.2C-D). Besides, trophoblasts exhibited the greatest expression changes among species compared to other placental cell classes, which was followed by stromal cells. With the evolutionary distance, the expression similarity in cells between humans and other mammalian species decreased, and this decrease occurred fastest in trophoblasts but at a similar rate in stromal cells, endothelial cells, and macrophages (Fig.2G). The rapid evolutionary divergence in trophoblasts likely represents a key driver for placental structural diversification across mammalian species.

### Evolutionary insights into trophoblast differentiation across mammalian species

To delve into the evolutionary drivers for trophoblast lineage diversification, we reconstructed differentiation trajectories of trophoblasts across multiple mammalian species. Previous research has proposed that the trophoblast subtypes are interrelated during development, with CTBs serving as the progenitors for both the EVTs through epithelium-mesenchymal transition (EMT) and invasion and STB via cell-cell fusion during placental development^40^. Specifically, we computationally ordered individual trophoblasts in primates (humans and macaca) in a 2D “pseudotime” trajectory to reconstruct their differentiation relationship (Fig.3A and Supplementary Fig.3A-B), following the method by Trapnell *et al*.^52^. In parallel, we constructed a trophoblast differentiation trajectory in rabbits, confirming that their CTBs differentiated into EVTs and STB similarly to primates (Fig.3B and Supplementary Fig.3C). In the rabbit EVT branch, known invasion-related genes (e.g., *SERPINE1*, *SERPINE2*, *RAC1*, and *DIO2*) were progressively upregulated during EVT differentiation, mirroring the expression pattern observed in humans (Fig.3C and Supplementary Fig.3D-F)^28, 40^. Additionally, we identified key regulators of the EVT differentiation path in rabbits, such as integrins beta-1 (*ITGB1*) and the metabolic enzyme *ALDH1A3* (Supplementary Fig.3E). Notably, *ALDH1A3*, which facilitates proliferation and invasion in murine glycogen trophoblast (GlyT) cells (analogous to human EVTs)^53, 54^, appeared to play similar roles in rabbit EVTs, suggesting conserved mechanisms for trophoblast invasion across species. Endogenous retroviruses (ERVs) have been proposed as a driving force for the evolution of mammalian placentas. *MFSD2A*, a receptor of the ERV envelope syncytin-2 (SYNC2) in human placentas^55^, was found to show increased expression along the rabbit STB fusion pathway. Also, the target gene *RAI14* of the ERV-derived enhancer MER50 was upregulated in the rabbit STB path, consistent with the findings in humans^56^ (Supplementary Fig.3D-E). Further, we identified multiple genes (e.g., *LRP8*, *GRB10*, and *TFEC*) as potential regulators in rabbit STB development, laying the groundwork for functional characterization in future studies (Fig.3C and Supplementary Fig.3E-F).

**Fig.3:**
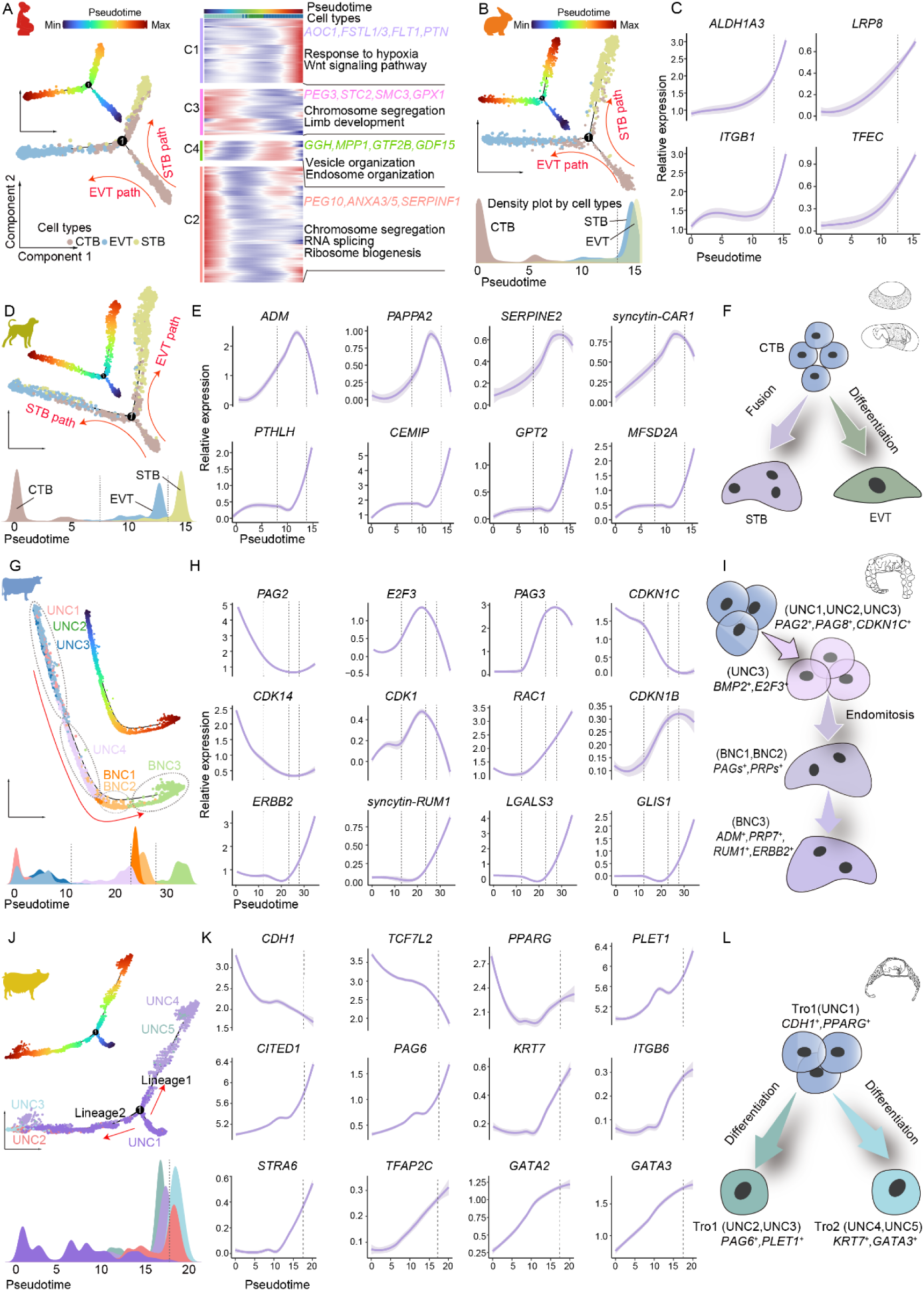
Reconstruction of the developmental relationships of mammalian trophoblasts through the pseudotime analysis. (A) The bivariate scatter plot illustrating the developmental pathways of human trophoblasts, diverging into EVTs and STB. The adjacent heatmap details dynamic changes in key biological processes during differentiation. (B-L) The developmental trajectories and gene expression patterns of trophoblasts from various placental types across pseudotime. Validation by Slingshot is documented in Supplementary Fig.3C, 4A, 4D, and 5C. (B, D, G, J): Single-cell trajectory analyses based on pseudotime showing the transformation of cell types from CTBs. Different colors denote various cell states or subgroups, illustrating the dynamic differentiation of placental trophoblasts. The density plots below highlight the distribution of cell types through pseudotime. (C, E, H, K): The key gene expression changes with pseudotime. Curves represent expression levels; purple shaded areas denote confidence intervals; grey dashed lines mark specific developmental timepoints aligned with the cell density plots. (F, I, L): Schematic diagrams of the molecular mechanisms underlying the differentiation of CTBs in discoid, zonary, cotyledonary, and diffuse placentas.

The subsequent comparative analysis of trophoblast differentiation trajectory in zonary placentas revealed a pattern parallel to that in discoid placentas (Fig.3D-F and Supplementary Fig.4A-B). We observed that starting from CTBs, one sub-branch highly expressed angiogenesis-(e.g., *ADM*, *PGF*, and *ANGPT2*) and trophoblast invasion-related genes (e.g., *PAPPA2*, *MMP2*, and *SERPINE2*), while the other exhibited pronounced expression of fusion-related genes. Both placental types displayed similar terminal differentiation features, characterized by activation of genes involved in vascular development, oxygen response, and Wnt signaling (Supplementary Fig.4B and Supplementary Data 4). In the STB differentiation trajectory, we observed conserved expression patterns shared by rabbits and humans, including upregulation of the transferrin receptor (*TFRC*) and ERV-related genes (*MFSD2A*) (Supplementary Fig.4C). Genes specific to canine STB differentiation included *CEMIP*, *PTHLH*, and *GPT2* (Fig.3E). Notably, our analysis revealed a distinctive expression pattern of *syncytin-CAR1*, a fusogenic ERV envelope gene conserved in Carnivora (Fig.3E). Unlike the conserved primate ERV genes *ERVW-1* (syncytin-1), *ERVFRD-1* (syncytin-2), and *ERVV-1*^57^, which were predominantly active along STB differentiation (Supplementary Fig.3D), *syncytin-CAR1* showed significant upregulation in both STB fusion and EVT differentiation, suggesting an unrecognized dual role in canine placental development (Fig.3E) and functional diversification of ERV-derived genes during mammalian placentation.

**Fig.4:**
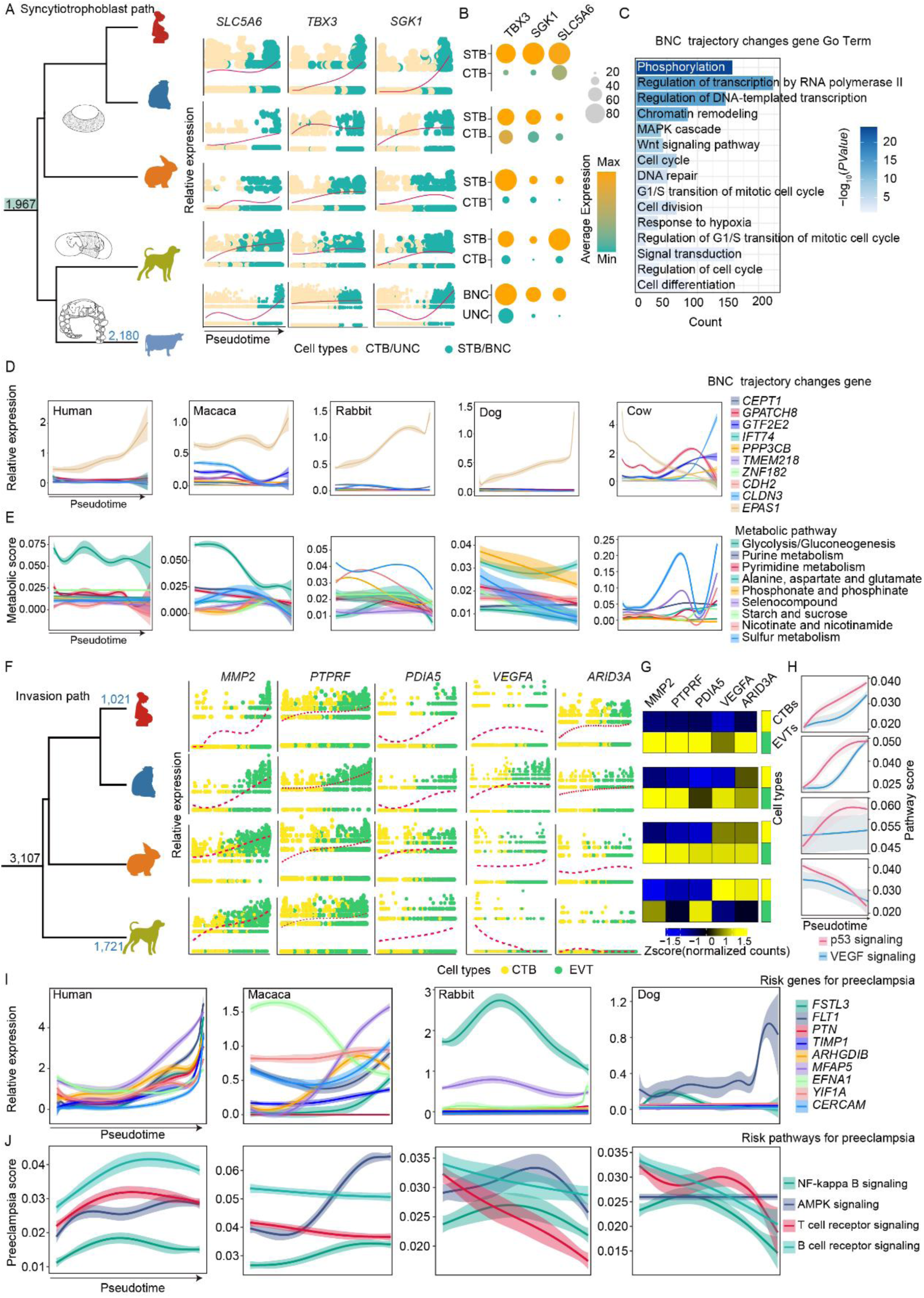
Conserved and divergent gene expression along trophoblast differentiation. (A) Evolutionary relationships among species and the number of conserved (black) and divergent (blue) gene expression trajectories in the fusion pathway. Branch lengths of the evolutionary tree have no biological meaning. The species from the top to the bottom are humans, macacas, rabbits, dogs, and cows. The middle section shows the expression changes of conserved and divergent genes along pseudotime during trophoblast fusion, with each row corresponding to a species and colors representing different cell types. (B) The dot plot showing the average expression levels of genes across different cell types (STB, CTBs, BNCs, and UNCs) in various species. (C) The bar chart showing the GO enrichment result of trajectory-changing genes in cows, with all enrichment results detailed in Supplementary Data 5. (D) The expression patterns of the exemplary genes along pseudotime in the trophoblast fusion pathways in different species (humans, macacas, rabbits, dogs, and cows). The expression trend in cows differs from that in other species. All related genes are listed in Supplementary Data 5. (E) Examples of the activity scores (AUCell) of conserved and divergent metabolic pathways during trophoblast fusion in cows. Detailed descriptions of metabolic pathways are provided in Supplementary Data 5, and activity scores for all pathways are shown in Supplementary Fig.8C-G. Of these, glycolysis / gluconeogenesis, phosphonate and phosphinate metabolism, and aminoacyl-tRNA biosynthesis were conserved metabolic pathways, while the others were pathways with increased specificity in cows. (F) Evolutionary relationships among species and the number of conserved (black) and divergent (blue) gene expression trajectories in the EVT pathway. The species from the top to the bottom are humans, macacas, rabbits, and dogs. The middle section shows the expression changes of conserved and divergent genes during pseudotime EVT differentiation, with each row corresponding to a species and colors representing different cell types. In the evolutionary tree, the black number (3,107) represents the number of genes with conserved expression patterns across all species in relation to humans (the alignment similarity ≥ 60%), while the blue numbers (1,721 and 1,021) represent the numbers of genes with trajectory changes in the EVT pathway (the alignment similarity ≤ 40%). All related genes are listed in Supplementary Data 5. (G) The heatmap showing the average gene expression levels across different cell types in various species. (H) The pathway scores of VEGF signaling and the p53 signaling pathway during EVT differentiation. The order of species from top to bottom is consistent with the evolutionary tree order in Fig.4F. (I and J) Examples of activity scores for reported PE-related genes (I) and pathways (J) during EVT differentiation across different species. Pathway activity scores for EVT differentiation across all species are shown in Supplementary Data 6.

Later, we reconstructed the differentiation relationships of cotyledonary placental trophoblasts (Fig.3G-I and Supplementary Fig.4D-E), which is consistent with our recent study at the spatial transcriptomic level^35^. The results showed that the pregnancy-associated glycoproteins (PAGs) and the prolactin-related proteins (PRPs) played crucial roles in the formation of BNCs in cows (Fig.3H and Supplementary Fig.4E). First, UNCs (start_UNC, UNC1, UNC2, and UNC3) with high expression of *PAG2*, *PAG8*, and *PAGE4* differentiated into another UNC subtype (mid_UNC, UNC4) that was characterized by high expression of *BMP2*, *E2F3*, *PEG10* as well as the cell cycle-related genes such as *BUB1B*, *SMC4*, and *CENPK*. Second, mid_UNC further differentiated into two BNC subtypes, with one BNC subtype (end1_BNC, BNC1 and BNC2) highly expressing PAGs (e.g., *PAG3*, *PAG14*, and *PAG18*) as well as PRP family genes (e.g., *PRP2*, *PRP3*, and *PRP14*) and the other (end2_BNC) highly expressing *PRP7*, *PRP6*, *HGF*, and *ADM*. Similarly, in primates, pregnancy-specific glycoproteins (PSGs) with analogous functions were found to participate in the formation of STB (Supplementary Fig.3A and D). In addition, we found substantial enrichment of key genes involved in cell cycle, DNA replication, chromosomal segregation, and cell division along the BNC pathway (Fig.3H and Supplementary Fig.4E), in sharp contrast to the human case that these genes were predominantly enriched in the EVT pathway and relatively sparse in the STB pathway^40^. Together with previous findings^46, 58^, our results demonstrate that BNC formation in ruminants relies on an endomitosis-rather than cell fusion-related mechanism, which is characterized by DNA replication and nuclear division but not cytokinesis^58^. At the initial stage of the transition from UNCs to BNCs, downregulation of *CDKN1C* (p57), *CDK14*, and *CDK1* suggests the resolution of cell cycle inhibition, thereby creating the necessary conditions for nuclear DNA replication^59^. At the terminal stage of differentiation, re-upregulation of cell cycle inhibitors, including *CDKN2A* (p16), *CDKN1B* (p27), and *CDK1*, facilitates the entry of mature BNCs into the state of cell cycle arrest, preventing excessive proliferation. Meanwhile, *CDK1* and *RAC1* were found to be upregulated during BNC formation, which, together with the previous study^60^, suggests that this upregulation might play critical roles in BNC formation by inhibiting cytokinesis. Additionally, cyclin-dependent kinases *CDK6* and *CDK8* were significantly upregulated at the late differentiation stage, likely to mediate the transcriptional regulation of pregnancy-specific proteins (PAGs and PRPs) to support the functional requirements of BNCs (Fig.3H and Supplementary Fig.4F). These dynamic gene expression changes reveal the highly coordinated interplay between cell cycle and differentiation during the UNC-to-BNC transition. Interestingly, in another species with cotyledonary placentas—goats, we also observed similar gene expression patterns, suggesting that BNC formation has a conserved role in ruminants (Supplementary Fig.4G). Furthermore, the *syncytin* gene (*RUM1*), which is of a retroviral origin and co-opted for placentation, along with the second phylogenetically unrelated *syncytin* gene, BERV K-1 (*Fematrin1*)^58^, was significantly upregulated during the endomitosis of BNCs. Notably, *syncytin-RUM1* was significantly enriched in mature BNCs (end2_BNC). At this stage, we also observed significant upregulation of invasive markers (e.g., *ERBB2* and *LGALS3*)^28^ and the reprogramming factor *GLIS1*^61^ (Fig.3H-I). The coordinated expression of these genes indicates that mature BNCs invade and eliminate the uterine epithelium (UE), and that with the influence of *syncytin-RUM1*, BNCs then fuse to form the syncytial plaques in place of the UE. These findings provide important molecular evidence for the formation of the unique structure at the maternal-fetal interface in the cotyledonary placentas.

Also, we reconstructed the trajectory of trophoblast differentiation in diffuse placentas (Fig.3J). We leveraged UNC1 with high differentiation potential (high expression of *CDH1*, *TCF7L2*, and *TP63*) predicted by cytoTRACE^62^ as the starting point of trophoblast differentiation (Supplementary Fig.5A-D), and found that trophoblasts developed along two differentiation trajectories (lineages 1 and 2), eventually becoming two different cell types (*PAG6*^+^ and *KRT7*^+^), which was further validated by slingshot^63^ (Fig.3K-L and Supplementary Fig.5C). This aligns with previous microscopic observations of two distinct types of trophoblasts with differential morphology^64^. No significant expression of cell fusion-related genes was detected, indicating that trophoblasts in diffuse placentas do not undergo cell fusion. Intriguingly, cells at the end of lineage 1 differentiation highly expressed genes encoding transport proteins associated with nutrition, metabolic regulation, and maternal-fetal material exchanges (*SLC45A3*, *SLC15A1*, *SLC4A7*, *SLC2A2*, and *SLC7A8*), sulfotransferases (*CHST8* and *CHST3*), and sialic acid transferases (*ST3GAL5* and *ST3GAL3*) (Supplementary Fig.5D-E), and these genes were also highly expressed in BNCs from cotyledonary placentas (Supplementary Fig.5F-G), suggesting that despite the absence of specialized differentiated trophoblasts (such as invasive or syncytial cells) during evolution, trophoblasts in diffuse placentas exhibit a similar function in maternal-fetal nutrient transport to the BNCs in cotyledonary placentas. Collectively, the differentiation pathways of the invasive trophoblasts were detected in discoid (primate and rodent) and zonary placentas, and syncytialization occurred in discoid, zonary, and cotyledonary placentas, while diffuse placentas had independent differentiation patterns, revealing significant differences and diversity during placental trophoblast differentiation in mammalian species.

**Fig.5:**
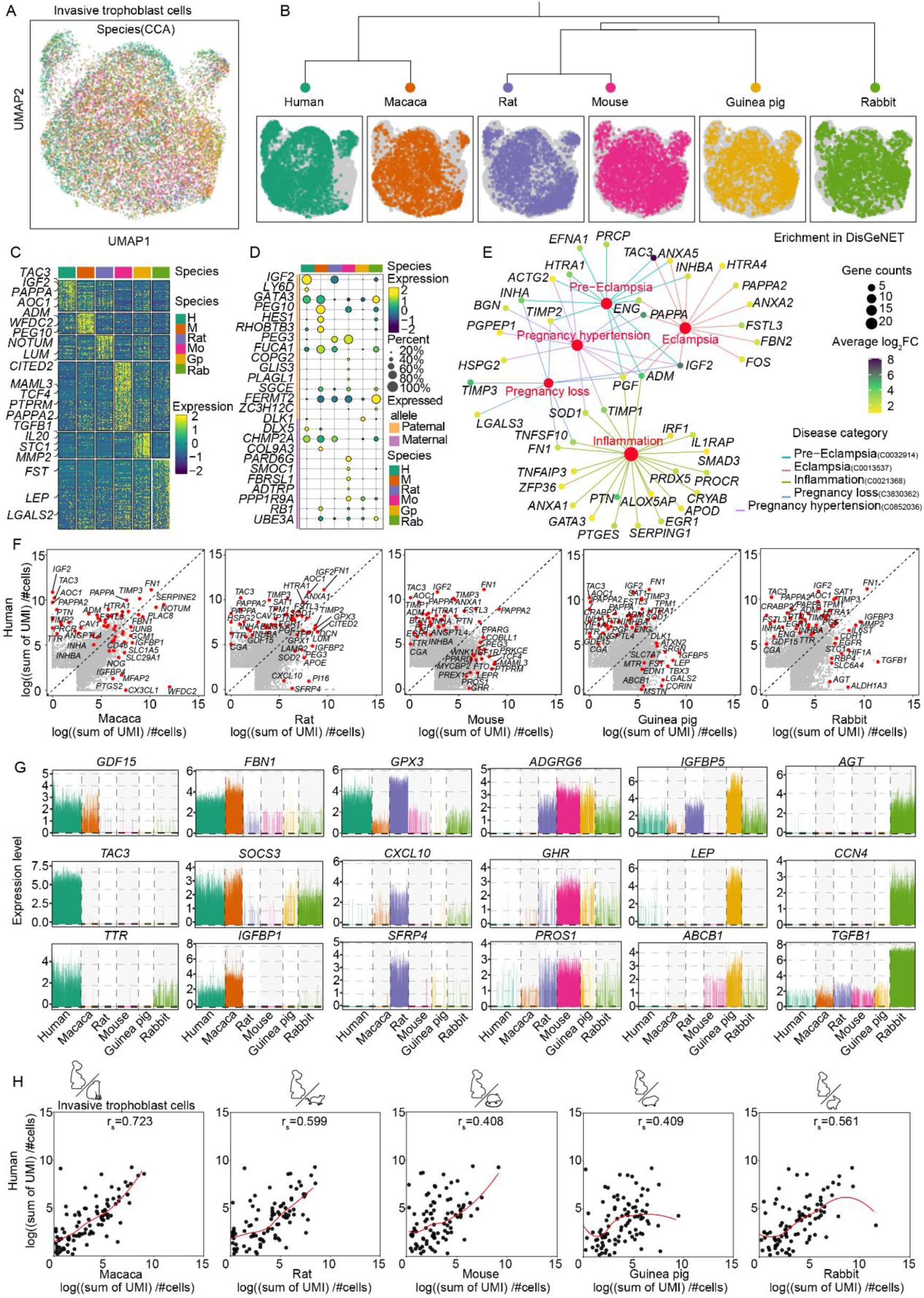
Evolutionary adaptation of human invasive trophoblasts to PE-related genes. (A) The UMAP plot of invasive trophoblasts from six species (humans, macacas, rats, mice, guinea pigs, and rabbits) integrated using the canonical correlation analysis (CCA), with different colors representing different species. (B) UMAP projections of invasive trophoblasts for each species, showing the clustering patterns of cells from each species. (C) The heatmap of differentially expressed genes (DEGs) across species, displaying normalized gene expression levels for each species. (D) The dot plot showing the expression levels of imprinted genes across species. The size of the dots represents the gene expression level, and the color corresponds to the species (Hu = humans, Ma = macacas, Rt = rats, Mo = mice, Gp = guinea pigs, Rab = rabbits). (E) Gene enrichment network displaying the enrichment of human DEGs in the DisGeNET database. The size of the gene nodes is related to the gene expression level, and the color indicates the association with specific disease categories. (F) The scatterplot showing the interspecies expression in EVTs. Gene expression is calculated as Log (sum of UMI) / number of cells. A subset of DEGs and markers are highlighted in red. (G) Expression patterns of selected PE-related genes across six species. (H) Correlation of PE-related genes (based on the intersection of DisGeNET and GeneCards databases with human DEGs, as detailed in Supplementary Data 8 in human EVTs with those in EVTs / invasive trophoblasts from other species.

### Conserved and divergent gene expression along trophoblast differentiation

Subsequently, we investigated the conserved and divergent genes and pathways involved in trophoblast differentiation by comparing the expression trajectories of one-to-one (1:1) orthologous genes across species. In brief, we used Genes2Genes (G2G)^65^ alignment to analyze the single-cell gene expression pseudotime trajectories between the reference and query species in pairwise comparisons, enabling us to track the conserved and divergent genes along similar differentiation trajectories (Supplementary Fig.6A-C and 7A-F). We identified 1,967 conserved genes involved in trophoblast fusion in primates, rodents, and *Laurasiatheria* (including dogs and cattle, Fig.4A and Supplementary Data 5), and found that the placental “core” gene *RALB*^6^ showed conservation in the STB pathway (Supplementary Fig.8A). In parallel, *TBX3*, a key regulator for CTB differentiation into STB^44^, was found to exhibit evolutionary conservation during trophoblast development. Additionally, genes such as *SLC5A6* (encoding a sodium-dependent multivitamin transporter), and *SGK1* (encoding a serine / threonine protein kinase) were conserved in trophoblast fusion (Fig.4A-B). Notably, *SGK1* has been reported to associate with pregnancy-related disorders including early pregnancy loss^66, 67^ and PE^68^ by affecting trophoblast invasion and endometrial decidualization. Here, our cross-species analysis revealed that *SGK1* might also play a critical role in linking trophoblast fusion / endomitosis to pregnancy complications. In cotyledonary placentas, unique BNCs develop within the chorion and fuse with the uterine surface epithelium to form syncytial plaques^35^. During BNC evolution, we identified 2,180 significant trajectory changes (Supplementary Data 5), primarily involved in processes such as cell division (GO:0051301), cell cycle (GO:0007049), phosphorylation (GO:0016310), chromatin remodeling (GO:0006338), and regulation of G1/S transition of mitotic cell cycle (GO:2000045) (Fig.4C). These biological processes, which are directly related to nuclear division, further provide crucial molecular foundations for the specialized BNCs in cotyledonary placentas that form through endomitosis of UNCs. Remarkably, significant expression changes were observed in *CEPT1*, *GPATCH8*, *GTF2E2*, *IFT74*, *PPP3CB*, *TMEM218*, and *ZNF182* during BNC evolution (Fig.4D), all of which are associated with placental morphology^6^. Similarly, the expression trajectories of *CDH2*, *CLDN3*, *ANXA2*, and the TF-encoding gene *EPAS1* also showed significant changes (Fig.4D and Supplementary Fig.8B). Of these, *CDH2* (encoding the N-cadherin) mediates calcium-dependent cell adhesion, crucial to maintenance of cell morphology and tissue structure^69^, while *CLDN3* (encoding the claudin 3) contributes to the formation of tight junctions, important to the intercellular barrier integrity^70^. Changes in these gene expression revealed the molecular basis for trophoblast specialization in cotyledonary placentas. To explore the evolution of placental nutrient allocation between the mother and the fetus, we further assessed the evolutionary changes in nutrient metabolism patterns during the fusion of STB and their impact on placental function (Fig.4E and Supplementary Fig.8C-G). We found that glycolysis / gluconeogenesis (hsa00010) showed conserved activity across all mammalian placentas, in alignment with previous human studies^71^ (Fig.4E). Similarly, pathways such as phosphonate and phosphinate metabolism (hsa00440) and aminoacyl-tRNA biosynthesis (hsa00970) exhibited conservation. In cotyledonary placentas, pathways such as starch-sucrose metabolism (hsa00500), alanine, aspartate and glutamate metabolism (hsa00250), and pyrimidine metabolism (hsa00240) displayed significant changes (Fig.4E), suggesting the specialized metabolic functions of BNCs in nutrient allocation.

**Fig.6:**
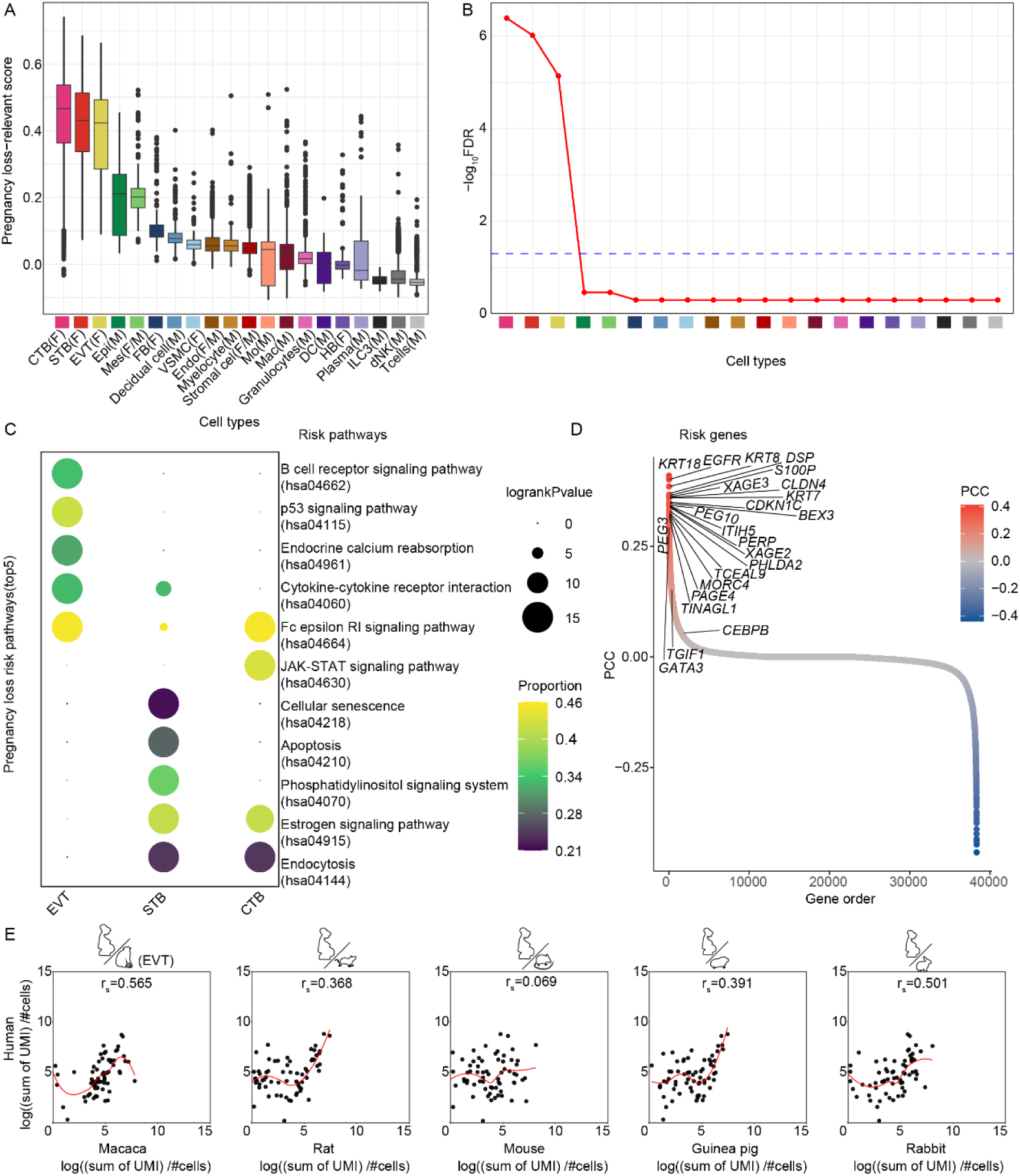
GWAS combined with single-cell transcriptomic analysis revealed the placental cell types associated with pregnancy loss. (A) Trait-relevant scores of pregnancy loss-related genes across various cell types. CTB, cytotrophoblasts; EVT, extravillous trophoblasts; STB, syncytiotrophoblasts; FB, fibroblasts; Endo, endothelial cells; Mac, macrophages; dNK, decidual natural killer cells; ILC, innate lymphocyte cells; VSMC, vascular smooth muscle cells; HB, Hofbauer cells; MES, mesenchymal cells; DC, dendritic cells; Epi, epithelial glandular cells. F and M in parentheses indicate the fetal and maternal origin, respectively. (B) The significance *P* values corrected by FDR showing the correlation between different cell types and pregnancy loss. (C) The dot plot showing the trait-relevant pathways identified by scPagwas for three cell types. The size of the dots represents the log-ranked *P* value for each pathway, and the color intensity reflects the proportion of cells influenced by genetic effects in each pathway (the pathway-level coefficient β > 0). (D) The trait-relevant genes ranked by the Pearson correlation coefficients (PCCs) using scPagwas across all individual cells. (E) Spearman correlations of the expression (UMI counts) of top 100 susceptibility genes of pregnancy loss (one-to-one (1:1) orthologous, Supplementary Data 8). From left to right are humans vs. macacas, humans vs. rats, humans vs. mice, humans vs. guinea pigs, and humans vs. rabbits.

Later, we assessed the conservation of gene expression trajectories in invasive trophoblasts across mammalian species, identifying 3,107 genes with conserved expression patterns along differentiation pathways of invasive trophoblasts (the alignment similarity ≥ 60%). These genes showed consistent conservation when subjected to the cross-species comparison with the human placenta (Fig.4F and Supplementary Data 5). For instance, *MMP2*, which degrades the endometrial extracellular matrix (ECM) to promote trophoblast invasion^72^, exhibited a conserved expression trajectory across placental types. Similarly, genes such as *CAPG* and *PDIA5*, which are involved in cell invasion^73^, exhibited conserved expression trajectories (Fig.4F-G and Supplementary Fig.9A). Within all mammalian invasive trophoblast differentiation, KEGG pathways related to invasion, energy metabolism, and angiogenesis, such as the HIF-1 (hsa04066), FoxO (hsa04068), and cell adhesion molecule (hsa04514) pathways, were steadily upregulated (Supplementary Fig.9B-C). Despite the conservation, we still identified approximately 1,723 trajectory changes during invasive trophoblast differentiation in zonary placentas in relation to humans (the alignment similarity ≤ 40%, Supplementary Data 5). These genes were enriched with pathways regulating the trophoblast invasion intensity, such as regulation of cell migration / growth, cell-cell adhesion, and cell-matrix adhesion (Supplementary Fig.9D and Supplementary Data 5). For instance, *VEGFA*, a key candidate in intravascular trophoblast invasion^74^, showed significant expression changes in zonary placentas (Fig.4F-G). Genes related to ECM degradation and remodeling, such as *CTSZ* and *ARID3A*^75^, also underwent changes in expression. Additionally, the VEGF signaling (hsa04370) and the p53 signaling pathway (hsa04115) were inhibited in EVTs in the zonary placentas (Fig.4H). It has been reported that inhibition of the VEGF signaling represses EVT proliferation, impairs their invasive capacity, and reduces angiogenesis^76^, and that p53 inhibition blocks decidualization while its activation promotes decidualization^77^. Hence, the current findings could help to address why trophoblasts in zonary placentas only invade the maternal endothelial tissue and exhibit limited decidualization. Surprisingly, during human EVT differentiation, the expression trajectory of genes (the alignment similarity ≤ 40%) and KEGG pathways associated with PE showed a reverse trend compared to that in other placental types (Fig.4I). To be more precise, genes such as *FSTL3* and *FLT1*, the important diagnostic markers for PE^78, 79, 80^, were specifically upregulated in human EVTs. Upregulated genes also included those involved in angiogenesis, hormone regulation, and tissue remodeling, such as *PTN*^81^, *TIMP1*^82^, *ARHGDIB*^83^, and *MFAP5*^84^ (Fig.4J). Consistently, pathways associated with PE, including NF-kappa B signaling (hsa04064)^85^, AMPK signaling (hsa04152)^86^, and T/B cell receptor signaling (hsa04660 and hsa04662)^87^, were specifically upregulated during human EVT differentiation (Fig.4K). Altogether, these findings reveal the molecular mechanisms underlying the diversification of placental structures during evolution and shed light on future basic and clinical explorations of pregnancy-related diseases.

### Evolutionary adaptation of human invasive trophoblasts to PE-related genes

Invasive EVTs are well-acknowledged as the key cell type involved in the pathogenesis of PE, with insufficient invasion being a key initiating factor for the pathogenesis of this placental disorder^88^. As described above, PE-related genes and KEGG pathways were upregulated in humans during EVT differentiation, while their expression was either downregulated or showed no significant change in other species. This observation prompted us to delve into the evolutionary expression patterns and the selective pressure on these genes in species with discoid placentas. To this end, we performed an integrative analysis of invasive trophoblast subtypes from all six species with discoid placentas (humans, macacas, rats, mice, guinea pigs, and rabbits). A total of 16,485 invasive trophoblasts were analyzed, including EVT cell types from humans, macacas, guinea pigs, and rabbits, the glycogen cells (GC) and trophoblast giant cells (TGCs) from mice that have been reported to exhibit invasion phenotypes similar to those in humans^54, 89, 90, 91, 92^, as well as invasive trophoblast subtypes in rats with significant expression of the prolactin superfamily (*PRL*) (Fig.5A-B)^91, 93^. The integrative analysis revealed that invasive trophoblasts were broadly distributed across species, with clustering patterns aligned to evolutionary relationships, i.e., cells from primate species (humans and macacas) were more similar in distribution, so as those with the rodent origin. Interestingly, while human cells overlapped with those from other species, they exhibited a relatively unique clustering pattern (Fig.5A-B), suggesting that human trophoblasts might have developed specific gene expression patterns or functional characteristics during evolution. Overall, we identified 2,365 conserved genes out of 11,068 orthologous genes (approximately 21%) across six species with discoid placentas, and these genes exhibited similar expression patterns across all species (Supplementary Fig.10A and Supplementary Data 7). Additionally, we identified 2,337 species-specific genes (approximately 21%) with significantly differential expression (|Log_2_Fold Change| ≥ 1, *P*_adj._ ≤ 0.05) (Fig.5C and Supplementary Data 7). Notably, the expression of these genes revealed the evolutionary characteristics of genomic imprinting in invasive trophoblasts. Specifically, 12 maternally imprinted genes showed species-specific expression in humans (*DLX5* and *HMP2A*), macacas (*COL9A3*), mice (*PARD6G*, *MOC1*, *BRSL1*, *DTRP*, and *PP1R9A*), and guinea pigs (*GRB10*, *PPP1R9A*, *RB1*, and *BE3A*), whereas 15 paternally imprinted genes displayed biased expression across the species with discoid placentas: *IGF2*, *LY6D*, and *GATA3* in humans, *PEG10*, *HES1*, *RHOBTB3*, and *IGF2R* in macacas, *PEG3*, *COPG2*, *GLIS3*, *PLAGL1*, and *SGCE* in mice, *FERMT2* and *ZC3H12C* in rabbits, and *DLK1* in guinea pigs (Fig.5D).

To delve into the evolutionary mechanisms for PE, we focused on 404 human-biased genes (|Log_2_Fold Change| ≥ 1, *P*_adj._ ≤ 0.05, and gene expression in over 20% of cells). These genes were significantly enriched with biological processes related to vascular development, VEGFA-VEGFR2 signaling, and ECM organization, all of which are closely associated with pregnancy-related diseases^94, 95, 96^ (Supplementary Fig.10B). Genes biased in primate (human and macaca) EVTs showed significant associations with PE (*P* < 0.01) (Fig.5E and Supplementary Data 7), and in humans, these genes were also highly associated with gestational hypertension and pregnancy loss (*P* < 0.01) (Supplementary Data 7), corroborating that PE specifically evolves in humans. Notably, the number of high-risk PE genes biased in human EVTs was one-to two-fold higher than that in invasive trophoblasts from the other five species with discoid placentas (Supplementary Fig.10C and Supplementary Data 7). Besides, through the analysis of the average pathogenic potential of these high-risk genes, we found that the levels of the highly pathogenic genes specifically expressed in human EVTs were over two-fold higher than those in invasive trophoblasts from the other five discoid placental species (Supplementary Fig.10D and Supplementary Data 7). These genes included those related to angiogenesis (*ADM*, *FSTL1*, *PGF*, *TAC3*, *CGA*, and *ANGPTL4*), members of the TGF-β family involved in immune response, cell migration, and proliferation regulation (*INHBA* and *INHA*), as well as those closely associated with trophoblast invasion (*FSTL3*, *PAPPA*, and *PAPPA2*) (Fig.5F and Supplementary Fig.10E). All these genes have been reported to be highly relevant to the critical roles of EVTs in remodeling the maternal spiral arteries and in invasion of the maternal decidua, and they have also been considered as key drivers for PE^97, 98, 99^. Besides, genes encoding proteins related to ECM and cell-cell junctions, such as *COL17A1*, *FN1*, *SPARC*, *TIMP3*, *TMEM59*, *ADAMTS1*, *VAMP8*, *COMMD6*, *PTGES*, and *ENG*, exhibited biased expression in human EVTs (Fig.5F, Supplementary Fig.10F-G, and Supplementary Data 7). These genes play critical roles in ECM degradation, tissue remodeling, and intercellular signal transduction regulation^99, 100, 101^. We also detected that PE-related genes were specifically highly expressed in EVTs from humans and macacas (Fig.5G). Illustrations of this point were *GDF-15* and *TAC3*, the potential biomarkers for PE^102, 103^, that were found to be highly expressed only in EVTs from the primate lineage, even after the inclusion of additional data from four mammalian species (cows, goats, pigs, and dogs) in our single-cell transcriptomic analysis (Supplementary Fig.10H). Similarly, *TTR*, a recently identified candidate biomarker for PE^104^, showed almost exclusive expression in human EVTs (Fig.5G). Yet, some genes associated with PE were also significantly expressed in invasive trophoblasts from rodent species, such as *ADGRG6*, *TGFB1*, and *PROS1* (Fig.5G), suggesting that less-invasive placentation models could still yield useful insights into the etiology of placental invasion and PE in humans. To gain more knowledge in this respect, we then compared the expression of PE-related genes in human EVTs with that in EVTs / invasive trophoblasts from other species, and found the highest correlation with macaca EVTs (r = 0.723), followed by rat invasive trophoblasts. In contrast, cells from mice, guinea pigs, and rabbits showed low to moderate correlations with those from humans in terms of the expression of PE-related genes (r = 0.406-0.561) (Fig.5H), suggesting that macacas and rats could be, in this sense, more fitting animal models for PE research. Collectively, these results reveal the evolutionary adaptation of PE-related genes in primate EVTs, providing novel insights into the etiology and molecular underpinnings of this disorder in humans.

### Linking pregnancy loss to specific placental cell types through the cross-omics analysis

GWAS is extensively leveraged to uncover underlying biological mechanisms and to predicate the disease risks by screening a large number of genetic variants across the genome to identify genetic loci associated with a specific phenotype or disease^105, 106^. Previous GWAS studies have revealed numerous genotype-phenotype associations related to pregnancy loss, but the mechanisms by which these genetic variants influence key biological pathways at the cellular level and drive disease progress remain enigmatic^18, 19^. Also, the significant association of biased genes in human EVTs with pregnancy loss (*P* < 0.01) (Supplementary Data 8) appealed to us. To acquire more knowledge in this regard, we collected the summary GWAS statistics from 114,761 pregnancy loss cases (including spontaneous abortion, missed abortion, and recurrent pregnancy loss) and 565,604 controls^18^ (Supplementary Data 8), as well as from approximately 140,000 human placental cells during the first^27^, second^29^, and third trimesters^40^ of pregnancy. With the pathway-based polygenic regression method (scPagwas)^107^, we integrated scRNA-seq with GWAS data and investigated the cellular context related to pregnancy loss in detail. Our results showed the significant association of EVTs, CTBs, and STB with pregnancy loss (FDR < 0.05) (Fig.6A-B and Supplementary Data 8), suggesting that maternal genetic effects play a crucial role in this disorder by regulating fetal trophoblasts. In EVTs, the pathways significantly associated with pregnancy loss included immune-related B cell receptor signaling (hsa04662), cell cycle-related p53 signaling (hsa04115), and the endocrine-regulated calcium reabsorption pathway (hsa04961) (Fig.6C). In STB, the significantly associated pathways chiefly involved cellular senescence and apoptosis, such as the cellular senescence pathway (hsa04218) and the apoptosis pathway (hsa04210) (Fig.6C). Of these, the cellular senescence^108, 109^ and JAK-STAT signaling pathways^110, 111^ have been identified to be closely associated with the risk of pregnancy loss. These findings suggest that diverse biological pathways are involved in pregnancy loss in different cell types. During human EVT differentiation, the pregnancy loss-associated B cell receptor signaling pathway showed upregulation, while in other species (e.g., macacas, rabbits, and dogs), it exhibited a downward expression trend (Supplementary Fig.11A), and during syncytialization, the pregnancy loss-related cellular senescence and apoptosis pathways progressively became active in all species except rabbits (Supplementary Fig.11B), suggestive of variations in pregnancy loss-associated pathways across different placental types and species during evolution. In addition, we identified high-priority genes related to pregnancy loss, including some expected ones (e.g., *KRT18*, *EGFR*, *KRT8*, *DSP*, *S100P*, and *CLDN4*) (Fig.6D) that were trophoblast-specific (Supplementary Fig.11C). Of these, *KRT18* plays a critical role in trophoblast migration and invasion, essential for embryo implantation^112^, while *EGFR* and *S100P* have repeatedly been reported to be implicated in miscarriage^113, 114^. Other interesting genes were also identified, such as *XAGE2* and *XAGE3* (members of the XAGE family of cancer / testis antigens^41^), *ITIH5*, and *BEX3* (a member of the Bex / Tceal family^115^). Next, we conducted a cross-species analysis for the expression of pregnancy loss-related genes, and found significant variability across species. For instance, *KRT18* and *KRT8* were significantly expressed in EVTs in humans, in STB in macacas, and in CTBs in rabbits and dogs, while *EGFR* and *DSP* also showed species-specific expression patterns across trophoblast subtypes (Supplementary Fig.11C). The further comparative analysis of invasive trophoblasts across species revealed that humans and macacas (Fig.6E) shared the highest gene expression similarity for pregnancy loss-related genes, followed by that between humans and rabbits (r = 0.565). Conversely, rats, mice, and guinea pigs exhibited significantly lower similarity (r = 0.069-0.391). These findings propose macacas and rabbits as promising candidate species for development of animal models to study pregnancy loss, particularly due to their genetic proximity to humans and the shared placental features.

In addition, we collected summary GWAS statistics from 750 cases of recurrent pregnancy loss and 150,215 controls^20^, and combined them with the corresponding human scRNA-seq data, employing the same means as described above. The results demonstrated that EVTs still exhibited significant association with recurrent pregnancy loss (FDR < 0.05). Mesenchymal stromal cells and macrophages also showed significant association (Supplementary Fig.12A-B). Key pathways involved in recurrent pregnancy loss included NF-kappa B and tight junction signaling, both related to inflammation and immune response, as well as pathways such as ECM-receptor interaction, the citrate cycle, and glycosaminoglycan biosynthesis, which also play important roles in pathogenesis of recurrent pregnancy loss (Supplementary Fig.12C). These recurrent pregnancy loss-associated pathways in EVTs exhibited evolutionary divergence and conservation during EVT differentiation. For example, NF-kappa B signaling was only detected in primates (humans and macacas) during differentiation, while tight junction signaling was downregulated exclusively in zonary placental species (e.g., dogs). In contrast, ECM-receptor interaction, the citric acid cycle, and glycosaminoglycan biosynthesis showed conserved activity in all species studied (humans, macacas, rabbits, and dogs) during EVT differentiation (Supplementary Fig.12D). The primary genes associated with recurrent pregnancy loss included several well-known ones, such as *FN1*, *JPT1*, *JPT2*, *RACK1*, *ITGB1*, *ELOB*, *ATF3*, and *FOS*, which are related to the growth, migration, and invasion of trophoblasts^116,103^ (Supplementary Fig.12E and Supplementary Data 8). Recent studies showed that the deficiency of *JPT2* could lead to the accumulation of citrate and reactive oxygen species (ROS), promoting macrophage polarization and therefore impairing the trophoblast functions, making it a potential therapeutic target for recurrent pregnancy loss^17^. We also identified additional genes associated with recurrent pregnancy loss, such as *ND4*, *ND5*, *ATP5F1A* (mitochondrial transcripts), and *ELOB* (Supplementary Fig.12E). In human EVTs, the expression of recurrent pregnancy loss-associated genes showed the highest correlation with that in macacas (r = 0.599) (Supplementary Fig.12F), whereas the correlation between humans and rats, mice, or guinea pigs was relatively low (r = 0.002-0.029). Taken together, we combined scRNA-seq data from all stages of human pregnancy with GWAS data of (recurrent) pregnancy loss, disclosing the key cell types (in particular trophoblasts) that are associated with (recurrent) pregnancy loss, as well as their roles in driving the disease.

### Key regulators of invasive trophoblasts and their potential roles in pregnancy disorders

To investigate key regulators driving invasive trophoblast diversification, particularly those influencing the biased expression of genes related to pregnancy diseases in humans, we employed SCENIC^117^ to evaluate the activity of orthologous regulators in invasive trophoblasts across six species (i.e., humans, macacas, rats, mice, guinea pigs, and rabbits), and defined a regulon specificity score (RSS) based on Jensen-Shannon divergence^118, 119^. We identified 186 regulatory factors with significant activity in the invasive trophoblasts from all species with discoid placentas, and found that while these factors retained essential physiological functions throughout evolution, their activity varied across species (Fig.7A and Supplementary Data 9). For instance, *TGIF1*, which encodes a protein pertaining to the three-amino acid loop extension (TALE) superclass of atypical homeodomains and playing a key role in regulation of the TGF-β signaling pathway^120^, showed high specificity in primates. *FOS* and *ATF4*, key regulators of cell proliferation, differentiation, and transformation^121^, also displayed significant species-specific regulatory activity in primates. In addition, *GATA3*, *CEBPB*, and *LHX4* exhibited unique regulatory activity in human invasive trophoblasts, whereas *RARB*, *ETV3*, and *HINFP*, the regulatory factors associated with germ layer differentiation, immunoglobulin gene activation, and vascular development^122, 123^, were specific to rodents.

**Fig.7:**
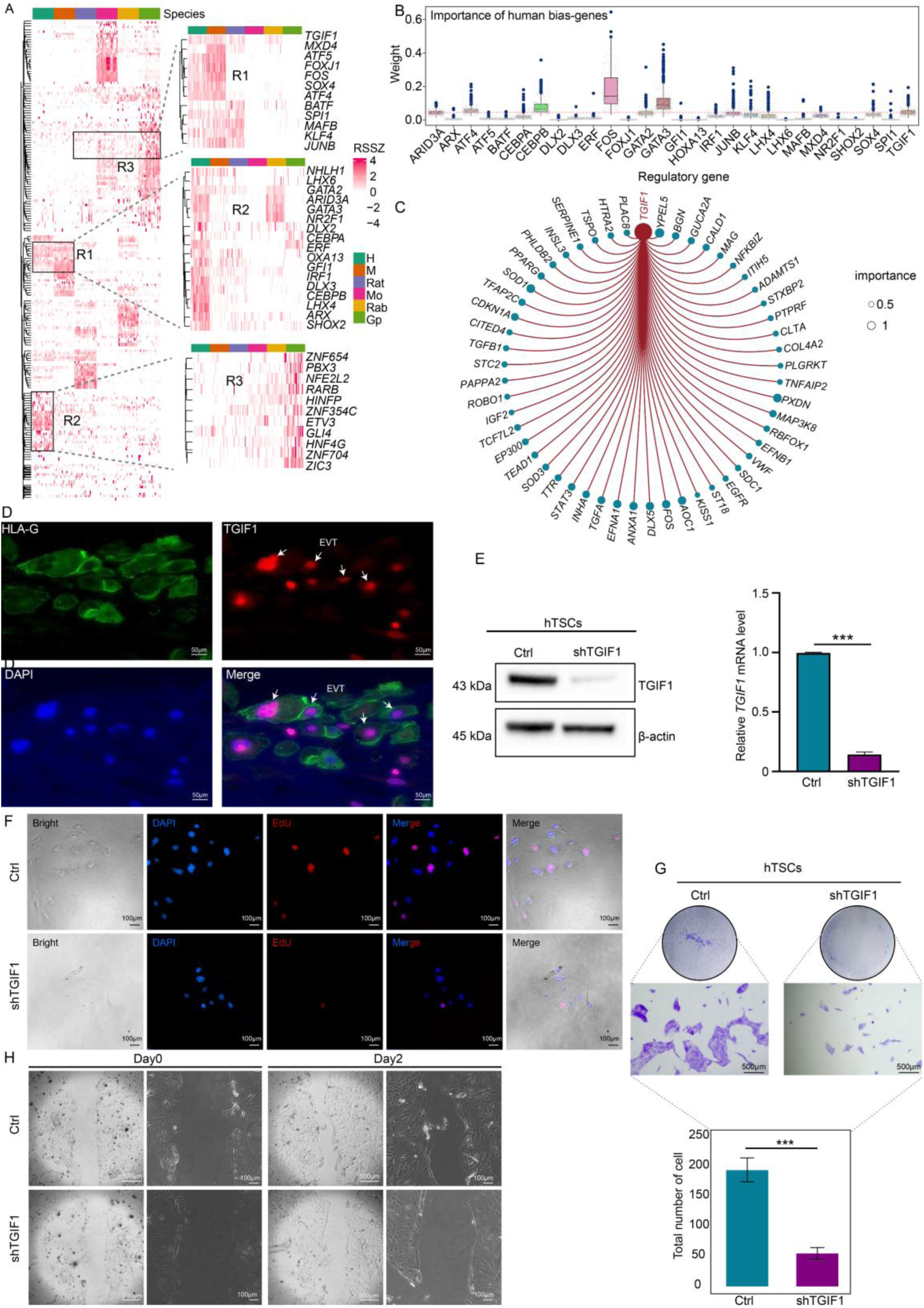
Key regulators of invasive trophoblasts and their potential roles in pregnancy disorders. (A) The heatmap displaying the activity of regulatory factors in invasive trophoblasts across six species. The regulatory factors are categorized into different regions (R1, R2, and R3) based on conservation, with black boxes highlighting the clusters of highly specific regulatory factors in primates and rodents. (B) The correlation between human-biased genes (avg_log_2_FC ≥ 1) and human-specific regulatory factors. (C) The regulatory network showing predicted *TGIF1* target genes in human EVTs. (D) The immunofluorescence analysis of human placental tissue (gestational week 40) showing TGIF1 staining (red) in the nuclei of HLA-G⁺ EVTs (green). Nuclei were counterstained with DAPI (blue). Arrows indicate EVTs co-stained with TGIF1 and HLA-G. (E) TGIF1 knockdown in hTSCs confirmed by Western blotting (left) and qRT-PCR (right). β-actin was used as a loading control. (F) The EdU incorporation assay. Red, blue, and magenta indicate the EdU⁺ cells, DAPI⁺ nuclei, and the merge. (G) The transwell invasion assay showing the impaired invasive ability of TGIF1-deficient hTSCs. Representative images (top) and quantification of invaded cells (bottom) are shown. (H) The migration assay showing the reduced migratory capacity in sh*TGIF1* hTSCs on days 0 and 2 post-scratch. Scale bars are indicated. Data are presented as the mean ± SEM of biological triplicates (n = 3). ***: *P* < 0.001.

To further elucidate human-specific gene regulatory networks, we applied GENIE3 (GeNeralized Influential Network Inference Engine), a machine learning-based network inference algorithm^124^, to the relevant single-cell transcriptomic data. The analysis revealed several key TFs that primarily drove the expression of human-biased genes, including well-known trophoblast TFs such as *GATA2/3*, *CEBPB*, *ATF4*, and *FOS* (Fig. 7B). These TFs have repeatedly been reported to be closely associated with pregnancy-related diseases such as PE and pregnancy loss^125, 126, 127, 128^, which is also consistent with the GWAS analysis (Supplementary Fig. 13A-C). Specifically, *TGIF1*, which ranked among the top six TFs in this study, drew our attention as its role in human or mouse placental trophoblast development has not been systematically explored. We subsequently analyzed the transcriptomic data and found that the *TGIF1* target genes in human EVTs not only included genes closely associated with pregnancy-related diseases such as PE and pregnancy loss, but also encompassed key genes involved in trophoblast development and invasive functions (Fig. 7C, Supplementary Data 10). Furthermore, *TGIF1* expression was found to be significantly higher in placentas from PE patients (Supplementary Fig. 13D).

Based on these findings, we then performed an immunofluorescence analysis on human term placental tissue (gestational week 40), and identified TGIF1 staining in the nuclei of HLA-G⁺ EVTs (Fig. 7D), suggesting its potential role in the EVT function. Subsequently, we knocked down *TGIF1* in human trophoblast stem cells (hTSCs) and the HTR-8/SVneo cell line (a human chorionic trophoblast cell line^129^) by introducing a *TGIF1*-shRNA expression vector into the cells via lentiviral transduction (Fig.7E, Supplementary Fig.13E), and observed markedly reduced trophoblast proliferation, invasion, migration, and clonogenic ability upon *TGIF1* downregulation (Figs.7F-I and Supplementary Fig.13F-J). Taken together, our results reveal that *TGIF1* contributes to the expression of pregnancy disorder-associated genes and regulates the invasive trophoblast function, providing insights into the molecular basis of conditions such as PE and pregnancy loss.

### TGIF1 target binding to trophoblast functional genes associated with PE

TFs exert their guiding role mostly by recognizing target DNA sequences to modulate chromatin and transcription. To delve into the mechanism by which *TGIF1* regulates trophoblast cell proliferation and migration, an RNA-seq analysis was performed on biological triplicates of control and sh*TGIF1* hTSCs (Supplementary Data 10). A total of 6,486 differentially expressed genes (DEGs, | Log2Fold Change | ≥ 1, *P*adj. ≤ 0.05) were identified, with the majority being downregulated in the sh*TGIF1* group (Fig. 8A). GO and KEGG enrichment analyses revealed that these downregulated genes were significantly enriched with several key pathways, including those related to cell proliferation (e.g., MAPK and PI3K-Akt signaling pathways), cell migration and invasion (e.g., ECM-receptor interaction and cell adhesion molecules [CAMs]), as well as immune response and cellular senescence (e.g., response to type I interferon, IL-17 signaling pathway, p53 signaling pathway, and cellular senescence) (Supplementary Fig.14 A and B). These findings are highly consistent with our phenotypic observations that knockdown of *TGIF1* impaired trophoblast cell proliferation, migration, and invasion.

**Fig.8:**
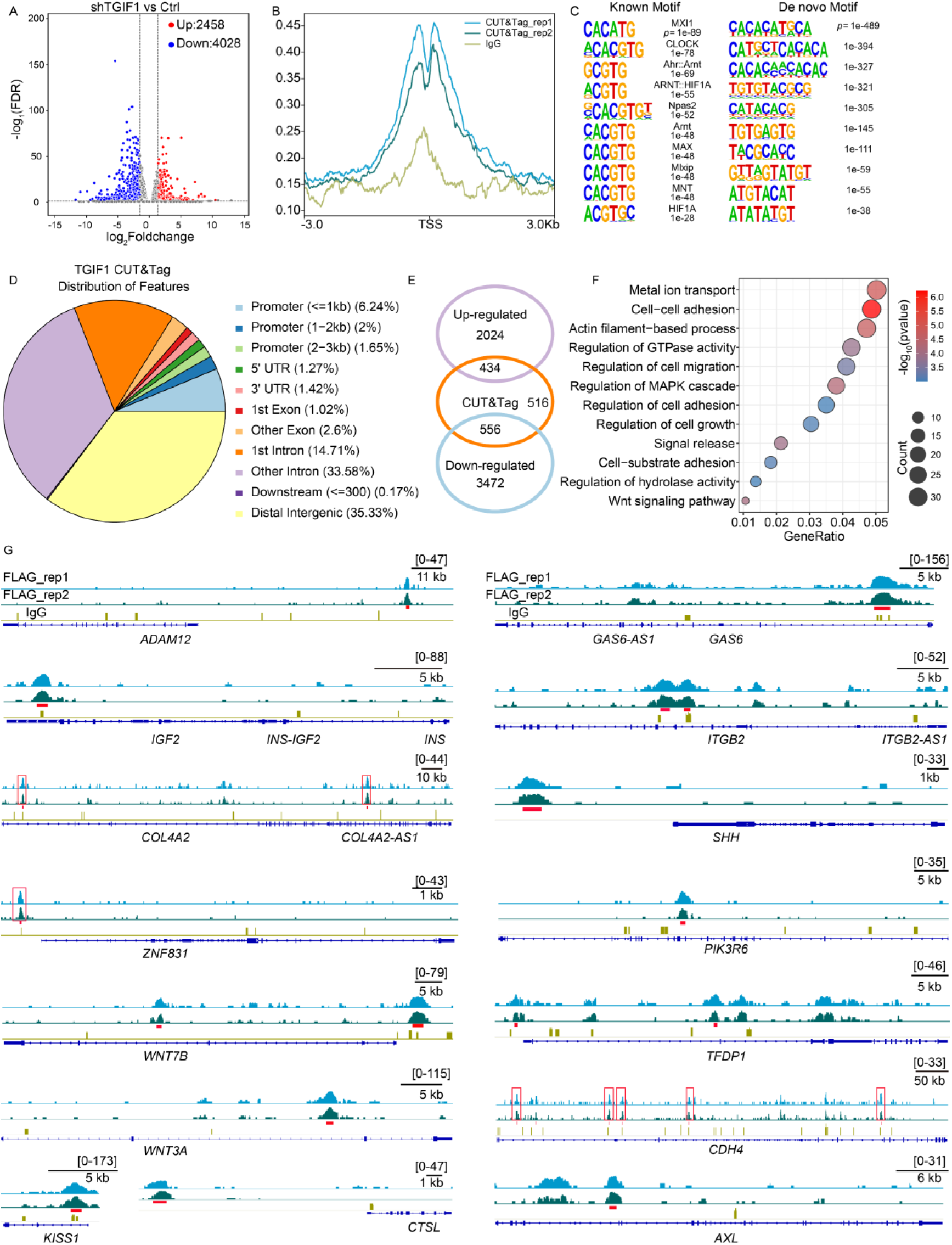
Genome-wide identification of the binding motifs of TGIF1. (A) The volcano plot of DEGs (sh*TGIF1* vs control). (B) Distribution of TGIF1 CUT&Tag signals across transcription start sites (TSS) (± 3 kb), showing promoter enrichment. (C) The DNA motif analysis of TGIF1 binding sites. Known motifs: previously reported motifs; *De novo* motifs: newly predicted motifs. (D) The genomic distribution profile of TGIF1 CUT&Tag peaks. (E) The Venn diagram showing the overlap between genes associated with TGIF1-binding sites and the DEGs identified by RNA-seq. (F) GO terms for the genes associated with TGIF1-binding sites. (G) Chromatin binding peaks of TGIF1 at genomic regions associated with pregnancy disorder-related genes.

Next, a CUT&Tag assay for TGIF1 was conducted on biological duplicates of hTSCs, with an aim to identify the TGIF1 DNA-binding sites in the human genome. In total, 3,528 to 4,012 peaks were identified, with an average width of 474.79 base pairs and an enrichment score of 5.18, corresponding to 3,108 to 3,481 annotated genes. By contrast, only 49 peaks were detected in the IgG control samples, indicating a high level of binding specificity (Supplementary Data 10). The density plots of TGIF1 DNA-binding sites reveal a broad spectrum of binding signals within gene body regions (Fig. 8B). The motif analysis of TGIF1 CUT&Tag peaks showed significant enrichment of multiple known and newly identified TF binding motifs (Fig. 8C, Supplementary Fig. 14C). A genome distribution analysis indicated that these peaks were predominantly enriched in intergenic regions, comprising 35.50% of all peaks. Interestingly, 9.89% of the peaks were located in putative promoter regions, 3 kb upstream of transcription start sites (TSS, Fig. 8D). Among the genes directly bound by TGIF1, 986 were differentially expressed between control and sh*TGIF1* groups, with 430 genes upregulated and 556 downregulated (Fig. 8E). The GO analysis revealed that these TGIF1 target genes were enriched with biological processes related to metal ion transport, cell-cell adhesion, regulation of GTPase activity, regulation of cell growth, and regulation of cell migration (Fig. 8F).

Specifically, Integrative Genomics Viewer (IGV) showed the TGIF1-binding peaks within the promoter or body regions of multiple genes closely associated with trophoblast cell proliferation, migration, invasion, and pathogenesis of PE and pregnancy loss (Fig. 8G, Supplementary Fig.14C). For example, *ADAM12* is a potential regulator of trophoblast invasion and a marker of healthy trophoblasts in the first trimester, with its expression consistently associated with the progression of PE and fetal growth restriction (FGR)^130^. *IGF2* plays significant roles in placental development, including promoting trophoblast proliferation, EVT migration, hormone secretion, glucose and amino acid uptake, and decreasing apoptosis^131^. *COL4A2*^132^, *KISS1*^133^, *SHH*^134^, *WNT3A*^135^, *RGL3*^136^, and *ZNF831*^136^ have been identified as risk-associated genes for human PE and recurrent miscarriage. In addition, *GAS6* signaling is capable of inducing PE, and inhibition of *AXL* prevents the disease progression in pregnant rats^137^. *PIK3R6*, *AXL*, and *WNT7B* are involved in regulation of the PI3K, TAM, and WNT signaling pathways, respectively, essential for trophoblast invasion and angiogenesis and strongly associated with PE^138, 139^. *PIK3R6* and *TFDP1* are also implicated in cell cycle regulation, potentially affecting trophoblast proliferation dynamics. Furthermore, several additional PE-related candidate genes, including *ITGB2*, *CTSL*, *EGFR*, and *DLX3*, also showed clear TGIF1 binding signals (Supplementary Fig.14E). These findings suggest that TGIF1 may contribute to the regulation of trophoblast function and the pathogenesis of PE through its interaction with *cis*-regulatory elements of a set of functionally critical genes.

## Discussion

The placenta plays indispensable roles in fetal development and maternal health, orchestrating key functions that support pregnancy and parturition. Unravelling the molecular features of placental cells, as well as their conserved and divergent patterns across different species, offers valuable insights into the mechanisms underlying the abnormal placental function. This knowledge could pave the way for novel diagnostic and therapeutic strategies for pregnancy-related complications such as PE and pregnancy loss. To date, high-throughput scRNA-seq has emerged as a powerful tool to construct comprehensive placental atlases in a variety of mammalian species including humans^27, 28, 40^, mice^89^, cattle^35^, and pigs^47^. However, these studies have been confined to a handful of species, leaving the full spectrum of placental diversity, as well as the molecular evolution of this important organ unaddressed. To fill in this gap, we presented here the first comprehensive single-cell transcriptomic atlases across ten representative mammalian species, representing the entire range of four placental types defined to date, namely the discoid, cotyledonary, diffuse, and zonary placentas. Our comparative analyses reveal significant evolutionary differences in trophoblasts and highlight their pivotal roles in the diversification of placental functions. In particular, we observed that trophoblasts evolved at a faster rate than other placental cell types, underscoring their central role in shaping species-specific placental morphology and functions. This rapid evolution of trophoblasts serves as a key driver in the molecular divergence of placental gene expression, laying the foundation for understanding trophoblast differentiation across species and the functional implications of this diversification.

To enhance placental efficiency and pregnancy health, placental and endometrial tissues undergo remodeling to minimize the maternal-fetal distance and maximize the interaction surface area as gestation progresses. This adaptation is observed in all mammalian species regardless of placental subtypes^64^. In cattle, the multicotyledonous structure exemplifies placental adaptation in a limited invasive context, primarily through the specialization of trophoblast giant BNCs^35^. This specialization is closely associated with the unique UNC differentiation into BNCs, a mechanism that remains largely hypothetical. Previous studies proposed the involvement of endoreplication or endomitosis in formation of polyploidy cells without direct evidence^46, 58^. Through comparative cross-species analyses, this study has elucidated, for the first time, the underlying mechanisms driving UNC differentiation into BNCs, highlighting the pivotal role of cell cycle regulators, where endoreplication initiation in UNCs is linked to inhibitory factors such as *CDKN1C* and *CDK1*, while BNC maturation depends on upregulation of *CDK1* and *RAC1* activity. This regulatory cascade is likely to suppress cytokinesis and excessive nuclear division, facilitating the formation of BNCs and illustrating the ubiquity of BNCs in ruminant placentas. Additionally, the metabolic specialization of BNCs further optimizes placental functions with significant enrichment in pathways such as starch-sucrose metabolism, alanine metabolism, and pyrimidine metabolism. This metabolic adaptation supports prolonged gestation and intricate nutrient exchanges in ruminants, significantly improving the functional efficiency of the placenta at the maternal-fetal interface. The functional adaptation of retroviral envelope genes in mammals has played a significant role in their evolutionary adaptability. In primates, *syncytin* genes are primarily associated with trophoblast fusion, whereas in zonary placentas, *syncytin-Car1* also participates in EVT invasion. In contrast, *syncytin-Ory1* in lagomorphs does not exhibit notable cell type-specific expression. However, in cotyledonary placentas, *syncytin-RUM1* is linked to endoreplication in UNCs, while *syncytin-RUM1* is enriched in mature BNCs. This gene specialization, along with the co-expression of invasion-related genes such as *ERBB2*, *LGALS3*, and *SERPINE1*, and the reprogramming factor *GLIS1*, is likely to form the molecular basis for the fusion of BNCs with uterine epithelial cells, further leading to the formation of syncytial plaques. These findings not only reveal the functional divergence among different species but also highlight the driving forces for trophoblast specialization and functional adaptive evolution, providing crucial insights into placental evolution.

The zonary (canine) placenta is characterized by restricted (shallow) invasion of trophoblasts, with no impact on the maternal capillaries and decidual cells. Thus, being structurally and functionally placed between noninvasive epitheliochorial placentation (cotyledonary and diffuse placentas) and the more invasive hemochorial type, it presents an interesting and important model to understand the evolutionarily determined factors in mammalian placentation^140^. Here, we found some similarities between zonary placentas and other invasive placental types (discoid placentas), indicating shared molecular mechanisms in different types of invasive placentas. We found many molecularly conserved features during the differentiation of their EVTs, e.g., *PAPPA2*, *MMP2*, and *MMP12* were highly expressed at the end of differentiation, supporting the invasive ability of EVTs on maternal tissues by regulating the degradation of ECM and the remodeling of the maternal-fetal interface^141, 142, 143^. This provides the molecular basis for use of the canine placenta as a model for translational and comparative studies on human placental pregnancy disorders. Despite the conservation of the trophoblast differentiation patterns between dogs and humans, specialization still occurred during canine invasive trophoblast differentiation. Specifically, canine EVTs showed a significant decrease in the expression of certain key molecules at the end of the differentiation pathway. An illustration of this point was *VFGFA*, a key candidate molecule involved in intravascular invasion, that maintained the high expression level in human EVTs to promote infiltration and remodeling of the maternal vasculature^74^. By contrast, in canine invasive trophoblasts, its expression sharply declined at the endpoint of differentiation, possibly responsible for the weaker or restricted intravascular invasion. Besides, the p53 signaling pathway was inhibited in canine EVTs during differentiation. As p53, a key regulator of apoptosis and proliferation, plays important roles in decidualization^77^, its inhibition might lead to the reduced or weakened decidualization in human placentas. Collectively, these molecular differences explain why the zonary placenta is less invasive compared to the discoid placenta, i.e., why the maternal vessels are not fully eroded and why extensive decidualization does not occur in the former^140, 144, 145^. The molecular differences also contribute to more profound knowledge about the roles of fetal trophoblasts and decidual cells in limiting invasion into maternal uterine structures, providing important clues for understandings of related pregnancy complications in humans.

The foremost hypotheses regarding the etiology of PE are the insufficient trophoblast invasion and placental ischemia induced by abnormal spiral artery remodeling, further resulting in the hypoxia stress and endothelial dysfunctions^104^. In this study, we observed that the expression of specific genes in invasive trophoblasts of primates was closely associated with angiogenesis, the insulin-like growth factor signaling pathway, and ECM remodeling. Enhanced trophoblast invasion in humans ensures efficient resource acquisition but on the other hand increases the risk of PE, reflecting an evolutionary trade-off. Besides, we identified the highest correlation in the expression of PE-associated genes between human invasive trophoblasts and their macaca counterparts, lending direct support to the use of primates as model animals for PE research. Nevertheless, the source of macacas is typically limited, and the cost of macaca rearing is quite high^146^. Intriguingly, we found that rats and rabbits showed a moderate correlation with humans in terms of the expression of PE-associated genes, such as *GPX3*, *CXCL10*, *SFRP4*, and *TGFB1*, providing alternative models to study PE and the related complications^145^.

By integrating GWAS data from pregnancy loss patients with single-cell transcriptomic data from human placental cells across pregnancy, we uncovered the placental cell types associated with (recurrent) pregnancy loss, as well as the critical genes and signaling pathways influencing pregnancy outcomes, such as *XAGE2/3*, *ITIH5*, and *BEX3*. Of these, *XAGE2/3*, a member of the XAGE family of cancer / testis antigens (CTAs)^41^, was found to be expressed in CTBs and STB. Because CTAs have been known as a large family of tumor-associated antigens expressed in various types of malignant tumors^147^ as well as ovarian and placental tissues^148^, associated with human tumorigenesis^147^, dysregulation of *XAGE2/3* is likely to interfere with the survival, invasion, and migration of trophoblasts, ultimately impairing embryo implantation and placental development, thus acting as a direct cause for pregnancy loss. *ITIH5* is a member of the serine protease inhibitor family^149^. It has been reported that the altered *ITIH5* expression is associated with tumorigenesis and metastasis^150^, and that *ITIH5*-mediated ECM stabilization suppresses invasion^149^. Based on these, we presume that *ITIH5* is associated with pregnancy loss probably by influencing the proliferation, migration, and invasion of trophoblasts. *BEX3* is a member of the Bex / Tceal family^115^, a multigenic cluster originating from a molecular domestication event involving transposable elements (TEs). This event, which occurs in the common ancestor of eutherian mammals, leads to the formation of a 14-gene cluster on the X chromosome in placental ancestors^115^. While many members in this family have yet been systematically investigated, previous studies have shown that the mammalian target of rapamycin (mTOR) pathway, which is related to a myriad of cellular processes such as cell growth, proliferation, and survival^151^, is one of the molecular networks in which Bex / Tceal genes are involved, with the roles of *BEX2* and *BEX4* in mTOR signaling uncovered^152, 153^. Therefore, it is plausible that dysregulation or mutation of the *BEX3* gene perturbs the trophoblast function, thereby increasing the risk of pregnancy loss. Collectively, the roles of *XAGE2/3*, *ITIH5*, and *BEX3* in embryonic and placental functions, and their potential contributions to pregnancy, warrant future investigation.

Besides, we found that the B-cell receptor and p53 signaling pathways in EVTs, as well as cellular senescence and apoptosis pathways in STB, were closely associated with the increased risk of (recurrent) pregnancy loss, suggesting the important roles of immune and apoptotic regulatory pathways in driving the progression of (recurrent) pregnancy loss. We also found that recurrent pregnancy loss was closely associated with placenta-derived macrophages, and that pathways such as allograft rejection, antigen processing and presentation, and growth hormone synthesis were strongly related to recurrent pregnancy loss. It has been known that while the embryo and the placenta are semi-allografts like transplanted organs, they can induce maternal tolerance and be free from a vigorous immune response^154^, which is the immunological foundation of successful pregnancy. Therefore, these pathways might influence the pregnancy outcomes by regulating immune tolerance at the maternal-fetal interface, and future studies are needed to elucidate the influence of macrophages on maternal-fetal immunologic tolerance to shield the embryo or the placenta from being rejected by the maternal immune system, eventually contributing to an elevated rate of successful pregnancy. In addition, our cross-species analysis revealed that macacas exhibited the highest similarity to humans in terms of the expression of risk genes for (recurrent) pregnancy loss, which was followed by rabbits, further suggesting the advantages of primates, as well as rabbits, a species with discoid placentas that has long been considered as a desirable model for studies on fetal growth and programming^155^, in modeling human pregnancy disorders, particularly in recapitulating the intricate pathogenesis of placentas.

Finally, we merged the risk genes for pregnancy loss and PE, as well as the regulatory networks in invasive trophoblasts across species, and identified several associated key TFs, such as *FOS*, *GATA3*, *CEBPB*, *GATA2*, and *TGIF1*. Previous studies have shown that *FOS*^125^, *GATA3*^126^, *CEBPB*^127^, and *GATA2*^128^ are directly associated with the risk of PE and pregnancy loss by influencing trophoblast migration and invasion. The biological significance of TGIF1 has been underscored in murine placental development through the use of knockout models. Specifically, the targeted deletion of *Tgif1* in mice resulted in reduced labyrinth vascularity and size, as well as diminished expression of the gap-junction protein Connexin 26, leading to placental insufficiency and severely growth-restricted embryos^156^. In human idiopathic FGR placentas, TGIF1 mRNA and protein levels were significantly elevated compared to those in controls. In BeWo cells, the knockdown of *TGIF1* via siRNA led to a significant downregulation of trophoblast differentiation markers at both the mRNA and protein levels^157^. These findings suggest that *TGIF1* acts as a potential upstream regulator of trophoblast differentiation, and that changes in *TGIF1* expression may contribute to abnormal villous trophoblast development. Our research indicates that TGIF1 directly regulates a group of PE risk genes by binding to their *cis*-regulatory elements, emphasizing its role as an upstream transcriptional regulator in the pathogenesis of PE.

In conclusion, by constructing a comprehensive single-cell transcriptomic atlas of placentas from ten mammalian species, this study has significantly advanced our understanding of the evolutionary conservation and divergence of placental trophoblasts, not only providing novel insights into the molecular basis of placental functions and dysfunctions, but also opening new avenues for development of improved models to study pregnancy disorders. Future research should focus on the functional validation of key regulatory genes and pathways identified in this study as well as their translational potential, with a long-term goal to augment pregnancy health outcomes.

## Methods

### Ethics statement

All experimental procedures and sampling protocols were reviewed and approved by the Institutional Animal Care and Use Committee (IACUC) of Northwest A&F University (Approval No: XN2024-0416). All sampling procedures strictly adhered to the “Guidelines for the Ethical Treatment of Laboratory Animals” established by the Ministry of Science and Technology of China. Detailed information on the samples, including strains, sources, rearing, ages, and genders, is available in Supplementary Data 1.

### Animals and tissue collection

In this study, maternal-fetal interface samples (including placentas, endometrium, and myometrium) were collected from multiple mammalian species, i.e., guinea pigs (*Cavia porcellus*), rabbits (*Oryctolagus cuniculus*), dogs (*Canis lupus familiaris*), cows (*Bos taurus*), goats (*Capra hircus*), and pigs (*Sus scrofa*). Detailed information on animals and sampling procedures is available in Supplementary Data 2.

### snRNA-seq library construction and sequencing

Single-nucleus RNA isolation was performed as previously described^158^. Placental tissues from different species were processed using the Chromium Single Cell 3’ GEM, Library & Gel Bead Kit v3 (PN-1000075) according to the manufacturer’s guidelines for library construction. After conversion using the MGI Easy Universal DNA Library Preparation Kit, the libraries were sequenced on a compatible BGISEQ-500 sequencing platform.

### snRNA-seq data processing

#### Raw sequencing data processing

The reads were paired-end, with Read 1 spanning 30 bases, where bases 1-20 corresponded to cell barcodes, and bases 21-30 were unique molecular identifier (UMI) sequences. Read 2 consisted of 100 bp of transcript sequences. The PISA software (https://github.com/shiquan/PISA) was employed to convert raw reads into FASTQ+ format according to the library structure, and to correct cell barcodes using the allow list when the Hamming distance was one or less. The reformed reads were aligned to the reference genomes using the STAR software^159^, and the specific reference genomes employed for each species were as follows: cows (*Bos taurus*, ARS-UCD2.0), goats (*Capra hircus*, ARS1.2), pigs (*Sus scrofa*, Sscrofa11.1), dogs (*Canis lupus familiaris*, UU_Cfam_GSD_1.0), rabbits (*Oryctolagus cuniculus*, UM_NZW_1.0), and guinea pigs (*Cavia porcellus*, mCavPor4.1). SAM files were converted into BAM files and annotated with the reference gene sets using the PISA software. UMIs from reads sharing the same cell barcode and gene annotation, even with a 1 bp mismatch, were adjusted to match the most frequent sequence. We then applied the DropletUtils R package^160^ to eliminate empty droplets. The final cell-by-gene matrix was constructed using PISA. We employed the Souporcell software^161^ to address potential contamination from ambient RNAs. To ensure cross-species consistency, all public datasets used in this study, including those from *Homo sapiens*^40^, *Cynomolgus macaque*^31^, *Mus musculus*^34^, and *Rattus norvegicus*^32^, were reprocessed from raw FASTQ files using the same PISA pipeline. All datasets underwent the same procedures as newly generated data. Cell annotations from the original publications were incorporated using the AddMetaData function in Seurat (v4.4.0). Markers reported previously were also used to verify cell identity assignments. For all species, we included all biological replicates at the selected developmental stages, except for the rat dataset, where one E19.5 sample (GSM8016898_19_5_7-PD) was excluded due to relatively low cell yield and reduced data quality (Supplementary Data 1).

#### Doublet filtering, elimination of the batch effect, and cell clustering

The final cell-gene matrix was imported into the Seurat (v4.4.0) package to create Seurat objects. Subsequently, normalization, scaling, and dimensionality reduction were carried out sequentially using the CreateSeuratObject, NormalizeData, FindVariableFeatures, ScaleData, and RunPCA functions, all with default parameters^48^. For quality control, cells were only retained if the number of detected genes was greater than 500 and the percentage of detected mitochondrial transcripts from MT genes was less than 20% (Supplementary Data 1). Variable genes were identified using the FindVariableGenes function in Seurat with default settings (selection.method = “vst”, nfeatures = 2000). The FindNeighbors (resolution = 1) and FindClusters functions were then employed to identify clusters based on principal components. DEGs between clusters were subsequently filtered with an adjusted *P* value < 0.01. Doublets were filtered using the Scrublet software^162^. All quality control, dimensionality reduction, clustering, and cell type annotation steps were performed independently for each species prior to cross-species integration.

### Inference of cell type identity

Cell type identification for each cluster was performed by plotting canonical cell markers using the FeaturePlot function in Seurat, and by identifying cluster-specific genes. Markers for each cell type were obtained from previous reports^27, 28, 31, 40, 46^, the CellMarker database^163^, and the Human Protein Atlas (https://www.proteinatlas.org). To validate the specificity and accuracy of cell type assignment, we conducted an overrepresentation analysis by comparing the top 200 cluster markers from our dataset with those from previously published human datasets^27, 40^ for the same cell types. To assess whether the overlap between two gene sets was significantly greater than what might occur by chance, we used the “enrichment” function from the R package bc3net (v1.0.4, https://github.com/cran/bc3net) that applies a Fisher’s exact test. The *P* values obtained from the Fisher’s exact test indicate the likelihood that the observed overlap between the two gene sets occurred randomly.

### Determination of the fetal or maternal origin

To interrogate the genetic origin of each barcoded cell without prior knowledge of individual genotypes, we employed the Souporcell tool implemented in Python^161^. The Souporcell tool infers the identity of cells by analyzing the aligned BAM files from each placental sample and by using the corresponding species’ genome files and filtered barcodes. For the post-Souporcell analysis, each cell was categorized based on its status (whether it was a singlet, doublet, or unassigned) and assigned an origin (either maternal or fetal, or classified as ambiguous). These identities were subsequently integrated into transcriptome-based dimensionality reduction data to further validate the maternal or fetal identity of each annotated cell type. Only cells that were confidently identified as singlets by the Souporcell pipeline were retained for further analyses.

### Identification of one-to-one orthologs and cross-species integration

As previously described^164^, one-to-one orthologs were identified using the EggNOG mapper (the β version) web interface and EggNOG (v.4.5) orthology data (http://eggnog5.embl.de)^165^. EggNOG functional terms were assigned to the predicted proteomes of humans (GRCh38.p14 assembly), macacas (T2T-MFA8v1.0 assembly), mice (GRCm39 assembly), rats (GRCr8 assembly), rabbits, dogs, cows, pigs, goats, and guinea pigs, focusing on one-to-one orthologs within mammalian species. Within each species, functional groups and genes were pruned based on the following criteria: (i) if a functional group mapped to multiple proteins from different genes; (ii) if different protein isoforms from the same gene mapped to distinct functional groups. The EggNOG functional group assigned to a gene was then used as an intermediate step to establish pairs. We only considered one-to-one orthologous pairs for further analyses, while the one-to-many or many-to-many orthologous pairs were discarded. After quality control and annotation within each species, we filtered the expression matrices to retain only one-to-one orthologs for cross-species integration. The canonical correlation analysis (CCA), as implemented in Seurat, was selected as the integration method as it outperformed other commonly used approaches including Harmony, BBKNN, fastMNN, scVI, and Scanorama, based on benchmarking using the scIB^166^ evaluation framework, achieving the overall best performance across batch correction and biological conservation metrics (Supplementary Data 1).

### DEGs, GO term and DisGeNET enrichment

In the global clustering, we performed the DEG analysis using the FindMarkers or FindAllMarkers function in Seurat. To analyze DEGs among different cell types, the FindAllMarkers function was applied. DEGs were defined as genes with the fold change ≥ 1 and adjusted *P* < 0.01. GO and DisGeNET enrichment analyses were performed using the compareCluster and enrichDGN function of clusterProfiler (v4.6.2)^167^. In the analysis, species-specific OrgDB packages were used for annotation: org.Bt.eg.db for cows, org.Ss.eg.db for pigs, org.Cf.eg.db for dogs, and org.Hs.eg.db for other species. Only GO and DisGeNET terms with Q < 0.05 were retained.

### Collection of TFs and risk genes for pregnancy diseases

All animal TFs were downloaded from AnimalTFDB^168^. A set of risk genes for pregnancy-related diseases were curated from the DisGeNET (https://www.disgenet.com)^169^ and GeneCards (https://www.genecards.org) databases. The pathogenic potential of these target genes was subsequently assessed using the VarElect analysis tool available within the GeneCards database. Imprinted genes were identified using resources from the Geneimprint database (https://www.geneimprint.com).

### Cell type comparisons across species

Following similar methods described in previous studies^170, 171^, we quantified the expression levels of each cell atlas as counts per million (CPM). To mitigate the impact of data sparsity that is common in low-coverage sequencing datasets, we used pseudo-cells to aggregate data from multiple cells of the same type. In this study, we selected 25 cells per pseudo-cell across all species. The use of pseudo-cells not only reduced computational costs but also enhanced expression estimation and clustering consistency. To systematically assess transcriptional similarities among cell types across different species, we performed an unsupervised MetaNeighbor analysis on the pseudo-cell data^172^. The MetaNeighbor approach is based on the premise that cells of the same type share more similar gene expression profiles than cells of different types. This method involves high variance gene (HVG) calculation, cell-to-cell correlation computation, cross-dataset validation, and neighbor voting. The average area under the receiver operating characteristic curve (AUROC) score was used to quantify the similarity between cell type pairs.

### The evolutionary variation of pseudobulk transcriptomes

For each species, we used the AverageExpression function in the Seurat R package to generate average (or pseudo-bulk) gene expression profiles for four major cell types: endothelial cells, trophoblasts, macrophages, and stromal cells. We then calculated the Spearman correlation between humans and other species using the cor function from the stats R package. To ensure comparability, nuclei were downsampled to an equal number for each donor within each subtype, which was repeated 100 times. In each iteration, the number of sampled cell nuclei varied within a range of 200 to 2,000. These Spearman correlations were subsequently visualized as scatter plots, with each species’ correlation compared to the evolutionary distance from humans, as determined by the Timetree website (https://timetree.org/). Similarly, we compared subtypes within species by calculating the average Spearman correlation across all pairwise comparisons of individuals.

### Differential gene expression across species

In line with previous recommendations^173^, we conducted the analysis beginning with the list of one-to-one orthologous genes across the compared species. The differential expression analysis for the pseudo-bulked count profiles of each cell type was performed using DESeq2 (version 3.40.0)^174^. To identify genes with significant differential expression between species pairs, we applied stringent criteria. Specifically, we set the FDR threshold to 0.001, along with the requirement that DEGs at least exhibited a two-fold change of expression (log_2_FC ≥ 1) that was detected in at least 20% of cells. To validate these DEGs, we utilized the FindAllMarkers and FindMarkers functions from the Seurat package, and conducted the differential expression analysis on the CCA-integrated objects. The identified species-biased genes by DESeq2 included all DEGs obtained by both the FindAllMarkers and FindMarkers functions. Based on this rigorous approach, we identified species-biased genes for each cell type in each species, specifically defining a biased gene as one that was significantly upregulated in that cell type relative to other species, as confirmed by both methods.

### TF analysis

The SCENIC analysis was carried out following the command-line protocol of SCENIC v0.11.0^117^. Seurat objects were downsampled to ensure an equal number of cells across all analyses and converted to the loom format using the “as.loom” function from the loomR v0.2.1.9000 package^175^. Subsequently, these loom files were imported into the Python environment using the “sc.read_loom” function from the Scanpy package. Gene regulatory networks (GRNs) were inferred using SCENIC, employing the GRBboost2 co-expression algorithm implemented in arboreto (v0.1.3) with default settings (v0.1.3), as previously described^176^. The UMAP based on the area under the curve (AUC) matrix was calculated through SCENIC. To quantify the specificity of regulons in human cell types, we calculated the Jensen-Shannon divergence (JSD)^118^ and RSS as previously described^171^. The Z-score normalized RSS (RSSZ) and AUROC scores were compared for each TF regulon. Regulons with the AUCell score greater than 0.1 and the RSSZ score greater than 1.0 were deemed significant. For non-human species, we employed the JSD based on TF expression profiles. Two normalized vectors representing TF expression levels in individual cells and cell type assignment were used to calculate the JSD between cell types. The TF specificity score (TFSS) was then derived to quantify the specificity of each TF across different cell types. We further generated a TFSS matrix, where rows and columns corresponded to TFs and cell types, respectively. TFs with the highest TFSS were considered essential for the respective cell types. The TFSS values were normalized to Z-scores (TFSSZ) for further analyses.

### Inference of cell trajectories

Slingshot^63^ and Monocle2^52^ were employed to infer cell trajectories for humans, macacas, rabbits, dogs, cows, and pigs. It has been proposed that in humans and macacas, these trophoblast subtypes are developmentally connected: CTBs serve as the progenitor compartment, generating EVTs through decidual invasion and STB through syncytium formation during placental development^27, 40^. Although similar developmental connections have not been reported in rabbits and dogs, these species exhibit trophoblast subtypes comparable to those found in humans. All trophoblast objects were extracted and downsampled to the same number of nuclei / cells (1,000), and the DDRTree method was used to reduce the dimension of cells (max_components = 2 and method = ‘DDRTree’). Then, the dimension reduction function was implemented to determine cell differentiation. Due to the unknown differentiation trajectory of porcine trophoblasts, we first employed CytoTRACE^62^ to assess cell differentiation. Subsequently, we used Monocle2 to construct the differentiation trajectory. Additionally, we cross-validated our findings by performing trajectory inference using the Slingshot algorithm. Finally, the ‘plot cell trajectory’ function was employed to visualize the differentiation trajectory of cells, while the ‘plot_pseudotime_heatmap’ function was used to generate heatmaps, and the ‘differentialGeneTest’ function was utilized to infer the genes involved in differentiation.

### The gene expression trajectory during trophoblast differentiation and fusion

To compare the conserved and divergent gene expression trajectories during trophoblast differentiation across humans, macacas, rabbits, dogs, and cows, we employed the Genes2Genes (G2G, v0.1.0)^65^ alignment framework. G2G is a dynamic programming (DP)-based alignment algorithm designed to align single-cell reference systems with query systems along any axis of progression (e.g., pseudotime). This method combines dynamic time warping (DTW) with gap modeling to capture both matches and mismatches between timepoints, optimizing the alignment of gene expression trajectories across species. In this study, we analyzed predefined gene sets: for the EVT differentiation pathway, 7,470 orthologous genes expressed in humans, macacas, rabbits, and dogs were used, whereas for the trophoblast fusion pathway, 7,405 orthologous genes expressed in humans, macacas, rabbits, dogs, and cows were used. Cells from each differentiation pathway were extracted from the Monocle2 analysis, and G2G was used to align the predefined genes between reference and query species at 15 interpolated timepoints. This framework enabled to identify lists of conserved and divergent genes during trophoblast differentiation across species, with the alignment similarity ≤ 40% defined as the most divergent genes. The specified genes were then visualized using the Monocle2 software to display their expression characteristics along the differentiation trajectory. For the analysis of KEGG pathway expression trajectories during trophoblast differentiation, we first retrieved all KEGG pathway information for humans using the KEGGREST (v1.38.0) R package and performed orthologous gene conversion. Pathways with fewer than three homologous genes were excluded. We then calculated activity scores for the retained KEGG pathways as cells progressed through differentiation. The AUCell^117^ analysis was conducted to assess pathway activity, based on the ranking of genes within the gene sets, by calculating the cumulative AUC. A higher AUC indicates more active expression of the gene set in the respective cell. Finally, we examined the changes in KEGG pathway expression across species, evaluating whether their activity increased or decreased during differentiation.

### Inference of gene and TF regulatory networks

We utilized GENIE3 (v 1.24.0)^124^ to infer the regulatory networks between TFs and genes from single-cell transcriptomic data. In this study, we input species-specific activated TFs along with the full set of genes towards humans or PE risk genes, to assess their correlation with the predicted expression of each target gene. The TF with the highest weight was designated as the most relevant regulator for the corresponding target gene expression.

### Enrichment of placental cell types in the GWAS signal of pregnancy loss

We obtained GWAS summary statistics for pregnancy loss that comprised 114,761 women who experienced pregnancy loss and 565,604 female controls from Iceland, Denmark, the United Kingdom, the United States, and Finland from deCODE genetics^18^ (https://www.decode.com). Cases were defined based on International Classification of Diseases (ICD) codes for spontaneous abortion, missed abortion, self-reported pregnancy loss (Supplementary Data 11), and recurrent pregnancy loss (750 cases and 150,215 controls) from the GWAS Catalog^19^. To obtain more precise results, we also collected single-cell transcriptomic data covering the entire stage of human pregnancy, including the first^27^, second^29^, and third trimesters^40^. Approximately 140,000 placental single-cell transcriptomes were used for subsequent analyses. The scPagwas^107^ software was employed to infer the cell types, related genes, and associated pathways linked to multiple consecutive miscarriages. Technically, scPagwas is a pathway-based polygenic regression method that integrates scRNA-seq data with GWAS data to identify crucial cellular contexts for complex diseases and traits. This method involves linear regression of pathway-activated GWAS signals derived from scRNA-seq data to identify a set of trait-associated genes, and these genes were then used to infer the most relevant cell subpopulations associated with the trait.

### Cell culture

The hTSCs were derived from human preimplantation embryos. Culture of hTSCs was performed as previously described^177, 178^. Briefly, hTSCs were maintained in the hTSC medium comprising DMEM/F12 (Gibco, 12634010), 0.2% fetal bovine serum (FBS, Gibco, A5669901), 0.1 mM 2-mercaptoethanol (Sigma-Aldrich), 0.5% penicillin-streptomycin (Sigma-Aldrich), 0.3% BSA (Sigma-Aldrich), 1% ITS-X supplement (Gibco), 1.5 mg/ml L-ascorbic acid (Sigma-Aldrich), 50 ng/ml EGF (PeproTech, 100-15), 2 μM CHIR99021 (Sigma-Aldrich, 860508P), 0.5 μM A83-01 (MedChemExpress, 909910-43-6), 1 μM SB431542 (Tocris), 0.8 mM valproic acid (Sigma-Aldrich), and 5 μM Y-27632 (Tocris). Cells were passaged every 4-5 days using TrypLE™ Express Enzyme (Gibco).

The immortalized human first trimester EVT cell line HTR-8/SVneo (HTR8)^129^ was obtained from the National Collection of Authenticated Cell Cultures. HTR8 cells were maintained in the RPMI 1640 medium (Invitrogen) supplemented with 10% FBS (Gibco, A5669901), 1% glutamine (Sigma-Aldrich), 1% non-essential amino acids (Sigma-Aldrich), 100 units / mL penicillin (Sigma-Aldrich), and 100 μM streptomycin (Sigma-Aldrich). Cells were cultured at 37 °C in an atmosphere of 5% CO_2_ in air, and passaged by light trypsinization before reaching confluency (no more than 20 consecutive passages).

### *TGIF1* knockdown in hTSCs and HTR-8/SVneo cells

The siRNA sequence targeting *TGIF1* (5’-CGGGATTGGCTGTATGAGCACCGTT-3’) and a scramble sequence (5’-CAACAAGATGAAGAGCACCAA-3’) were validated and cloned into the shRNA expression vector pGreenPuro (System Biosciences), following the manufacturer’s instructions. The cloned shRNA constructs were verified through Sanger-sequencing. To produce lentiviruses, the lentiviral backbone plasmids (pMD2.G), packaging plasmids (psPAX2), and the constructed transfer plasmids (pGreenPuro) were co-transfected into HEK293T cells using a liposome-based transfection reagent (GoldenTran). Around 16 h post-transfection, the cells were refreshed and cultured for an additional 48 h. The supernatant was then collected and concentrated using the lentivirus concentration reagent (Biodragon), according to the manufacturer’s instructions. The harvested and concentrated viruses were subsequently used to infect hTSCs and HTR-8/SVneo cells in the presence of polybrene. Puromycin was adopted to select successfully transduced cells, as the shTGIF1 vector carried both a puromycin resistance gene (PuroR) and a GFP reporter.

### Total RNA extraction and quantitative real-time PCR (qPCR)

Total RNAs were isolated using a RNeasy Mini Kit (Qiagen). cDNAs were synthesized using the iScript Reverse Transcription Supermix kit (Bio-Rad) and amplified with SYBR Green PCR Master Mix (YEASEN) on a Touch Thermal Cycler Real-Time PCR system (Roche, LightCycler480). *GAPDH* was used as an endogenous reference gene. The relative changes in gene expression were calculated using the 2^-ΔΔCt^ means. The primers used in this study were as follows:

*GAPDH*-F, AGATCCCTCAAATGAGCTGG

*GAPDH*-R, GGCAGAGATGATGACCCTTTT

*TGIF1*-F, ACAAGGCTTCCTCAGTGGT

*TGIF1*-R, CTTGGGCTGTGAATGTGGAAG

### RNA-seq analysis

Raw sequencing reads were processed and filtered using software fastp with default parameters. The high-quality reads were then aligned to the human reference genome GRCh38.p14 using STAR with default parameters. FeatureCounts were then used to summarize the reads mapped to each gene. To avoid confounding effects from low-quality gene models, only high-confidence gene models with an FPKM of more than 1 in at least one sample were considered expressed.

### CUT&Tag assay and data analysis

The CUT&Tag assay was conducted by Wuhan Zhenyue Biotechnology Co., Ltd as previously described^179^. Cells were bound to concanavalin A-coated magnetic beads, and incubated overnight at 4 °C with the primary antibody rabbit anti-TGIF (1: 50; ab52955, Abcam) or IgG control (1: 50; RA1008-01, Vazyme). CUT&Tag was used to construct sequencing libraries following the protocol provided with the Hyperactive Universal CUT&Tag Assay Kit for Illumina Pro (TD904, Vazyme Biotech Co., Ltd). Trimmomatic (v0.39) was used to filter out low-quality reads. Clean reads were mapped to the human reference genome (GRCh38.p14) by Bowtie2 (v2.4.4). Samtools (v1.13) was used to remove potential PCR duplicates. MACS2 software (v2.1.1) was used to call peaks by default parameters. If the midpoint of a peak is located closest to the TSS of one gene, the peak is assigned to that gene. HOMER (v3) was used to predict motif occurrence within peaks with default settings for a maximum motif length of 12 base pairs. The target peak was visualized using IGV (v2.15.2). The GO enrichment analysis was performed using the Metascape^180^ analysis toolkit, and terms with *P* values less than 0.05 were considered significant.

### Cell proliferation and clonogenic assay

An *In Vitro* Imaging Kit (RiboBio, Guangzhou, China) for the detection of 5-ethynyl-2′-deoxyuridine (EdU) incorporation was used according to the manufacturer’s protocol. The EdU-labeled cells were imaged with a confocal microscope. Also, we utilized a CCK-8 kit (Beyotime, China) to evaluate cell proliferation via the WST-8 reagent. In brief, cells were first seeded into 96-well plates and incubated at 37 °C in an atmosphere of 5% CO_2_ in air. Following 24, 48, 60, and 72 h of incubation, 10 μL CCK-8 solution was added to each well, and cells were incubated for an additional 2 h. Absorbance was then measured at 450 nm using a microplate reader (Multiskan; Thermo Fisher Scientific) to assess cell viability and proliferation.

For the clonogenic assay, 100 cells were plated in 12-well plates and cultured under standard conditions (37 °C, 5% CO₂) for 7 days to allow colony formation. Later, the cells were gently washed with PBS, fixed with 4% paraformaldehyde (PFA) for 15 min at room temperature, and stained with 0.1% crystal violet solution for 30 min. Excessive staining was removed by rinsing with distilled water. Images of the colonies were captured under a microscope.

### Western blotting

Whole-cell lysates were prepared using RIPA lysis buffer (Solarbio, China), and protein concentrations were measured with the BCA Protein Assay Kit (Solarbio, China). Equal amounts of total proteins (15 μg per sample) were separated by 10% SDS-PAGE and transferred onto PVDF membranes (Millipore). Membranes were blocked with 5% BSA in TBST (Tris-buffered saline with 0.1% Tween-20) for 1 h at room temperature, followed by overnight incubation at 4 °C with primary antibodies mouse anti-TGIF (1: 1000; Santa Cruz, sc-17800) and rabbit anti-β-actin (1: 1000; Cell Signaling Technology, #4970). After washing, membranes were incubated with the corresponding HRP-conjugated secondary antibodies (Jackson ImmunoResearch, USA) for 1 h at room temperature. Protein bands were detected using enhanced chemiluminescence (ECL) reagents (Epizyme, China) and visualized using a ChemiDoc imaging system (Bio-Rad).

### Immunofluorescence staining

Immunofluorescence staining was performed on rehydrated tissue slides. Term placental villus FFPE sections (40 weeks of gestation) were provided by the corresponding author, Dr. Zhenyu Xiao. Antigen retrieval was conducted in 0.01 M sodium citrate-HCl buffer at 95 °C for 10 min, followed by natural cooling to room temperature. Slides were then blocked with 2% BSA in PBS for 30 min and incubated overnight at 4 °C with primary antibodies mouse anti-HLA-G (1: 1000; Santa Cruz, sc-21799) and rabbit anti-TGIF (1: 1000; Abcam, ab52955). After three washes with PBST, slides were incubated with the corresponding fluorescent secondary antibodies for 30 min at room temperature. After washing, sections were counterstained with DAPI and mounted using VECTASHIELD Antifade Mounting Medium. Images were acquired using a ZEISS LSM 880 confocal microscope and processed with ZEN (black) software.

### Transwell invasion and migration assays

For the invasion assay, 50 μL diluted Matrigel glue was added to the upper transwell chamber and placed in an incubator at 37 °C for 30 min. Then, transfected cells were adjusted at the density of 2 × 10^4^ / mL, and 100 µL cell suspension was added to each well of the upper chamber, with 500 µL DMEM containing 10% FBS added to the lower chamber. After cell adherence, the serum-free culture medium was changed and the cells were incubated for 12 h at 37 °C in an atmosphere of 5% CO_2_ in air. The cells were then moved, and the culture medium was discarded. The cells in the upper chamber were gently wiped, fixed with 4% PFA for 10 min, stained with 0.1% crystal violet for 10 min, and observed under an optical microscope. The number of invaded cells was subsequently counted.

For the migration assay, cells were seeded to 6-well plates at the density of 5 × 10^5^ cells per well and incubated for 12 h. Once the cells reached approximately 95% of confluency, a 200 μL pipette tip was used to create scratches across the monolayer. Images of the scratched areas were captured at 0 and 24 h (HTR-8/SVneo) or 48 h (hTSCs) using an optical microscope (Olympus). The scratch width at 24 or 48 h was measured and compared to the baseline using the Image-Pro Plus 6.0 software. The experiment was repeated three times, with each harboring three biological replicates.

## Data availability

The raw and processed data generated in this study have been deposited in the NCBI database under the accession code PRJNA1177647 (https://dataview.ncbi.nlm.nih.gov/object/PRJNA1177647?reviewer=jsgtf0iutq4ecm439a8 1pgh6k8). Human placental scRNA-seq data used in this study were obtained from the European Genome-phenome Archive (EGA, https://www.ebi.ac.uk/ega/; accession EGAS00001002449). Mouse placental snRNA-seq data were retrieved from GEO (accession: GSE156125). Macaca placental datasets were obtained from GEO (accessions: GSE180637), and the rat scRNA-seq data were also sourced from GEO (accession: GSE206086). GWAS summary statistics for pregnancy loss were downloaded from deCODE (https://www.decode.com/summarydata/), and those for multiple consecutive miscarriages were obtained from Estonian Biobank (http://www.geenivaramu.ee/tools/misc_sumstats.zip). All other relevant data supporting the key findings of this study are provided in the article or as Supplementary Data.

## Code availability

This paper does not report original codes. Any additional information needed to reanalyze the data reported in this paper is available from the lead contact upon request.

## Acknowledgements

This work was supported by the National Key R&D Program of China (Grant No. 2022YFF1000100 to Y.J.), the Key R&D Program of Shaanxi Province (Grant No. 2024NC-YBXM-096 to Y.Z.), the National Natural Science Foundation of China (Grant No. 82371685) and Beijing Nova Program (to Z.X.), and the earmarked fund for CARS-35-PIG. We thank the High-Performance Computing Platform of Northwest A&F University and the Computing Center in Xi’an for provision of computing resources.

## Contributions

Conceptualization, Y.J., Y.Z., Z.X., and X.W.; Methodology, G.T., A.Z., X.C., J.Y., Y.C., F.W., and T.S.; Investigation, G.T. (major), A.Z. (major), X.C., J.Y., Y.C., F.W., T.S., H.L., M.W., W.H., H.S., J.R., and F.L.; Writing – Original Draft, G.T., A.Z.; Writing – review & editing, all authors; Funding acquisition, Y.J., Y.Z., Z.X., and X.W.; Resources, Y.J., Y.Z., Z.X., and X.W.; Supervision, Y.J., Y.Z., Z.X., X.W., F.W., and T.S.

